# Inhibition of histone readers bromodomain extra-terminal proteins alleviates skin fibrosis in experimental models of scleroderma

**DOI:** 10.1101/2020.08.07.242198

**Authors:** Sirapa Vichaikul, Mikel Gurrea-Rubio, M. Asif Amin, Phillip L. Campbell, Qi Wu, Megan N. Mattichak, William D. Brodie, Pamela J. Palisoc, Mustafa Ali, Sei Muraoka, Jeffrey H. Ruth, Ellen N. Model, Dallas M. Rohraff, Jonatan L. Hervoso, Yang Mao-Draayer, David A. Fox, Dinesh Khanna, Amr H. Sawalha, Pei-Suen Tsou

## Abstract

Binding of the bromodomain and extra-terminal domain proteins (BETs) to acetylated histone residues is critical for gene transcription. This study sought to determine the anti-fibrotic efficacy and potential mechanisms of BET inhibition in systemic sclerosis (SSc). Blockade of BETs was done using a pan BET inhibitor JQ1, BRD2 inhibitor BIC1, or BRD4 inhibitors AZD5153 or ARV825. BET inhibition, specifically BRD4 blockade, showed anti-fibrotic effects in an animal model of scleroderma and in patient-derived diffuse cutaneous (dc)SSc fibroblasts. Transcriptome analysis of JQ1-treated dcSSc fibroblasts revealed differentially expressed genes related to extracellular matrix, cell cycle, and calcium signaling. The anti-fibrotic effect of BRD4 inhibition was at least in part mediated by downregulation of Ca^2+/^calmodulin-dependent protein kinase II α (CaMKII-α) and reduction of intracellular calcium concentrations. These results suggest that targeting calcium pathways or BRD4 might be novel therapeutic approaches for progressive tissue fibrosis.

## Introduction

Systemic sclerosis (scleroderma, SSc) is an autoimmune disease characterized by vascular dysfunction as well as excessive synthesis and deposition of extracellular matrix (ECM) in affected organs. Activation of the immune system and vasculopathy precede fibrosis, in which fibroblast activation and subsequent myofibroblast transdifferentiation are necessary events. At present, SSc is not curable; current treatments focus on managing disease manifestations in an effort to ease the progression of tissue fibrosis.

Although the exact etiology of the disease is not known, a growing body of literature has been pointing to the critical involvement of epigenetic mechanisms in SSc pathogenesis (1). Epigenetic regulation affects chromatin dynamics and thereby modulates gene transcription. In addition to DNA methylation and non-coding RNAs, histone changes are implicated in SSc fibroblast activation (2–4). Acetylation of histones on lysine residues is one of the most common histone modifications that relaxes the chromatin structure by loosening the histone-DNA interaction, which results in increased chromatin accessibility for transcription. Dysregulation of histone acetylation could result in aberrant gene expression leading to pathogenic consequences. These histone marks are therefore tightly regulated by a set of histone acetyltransferases and histone deacetylases. They are also controlled by proteins containing the bromodomain module, such as the bromodomain extra-terminal domain (BET) family proteins. The four members of the BETs, BRD2, BRD3, BRD4, and the testis-specific BRD-t, share a common domain consisting of two N-terminal bromodomains, BD1 and BD2, that bind to acetylated lysine residues on histones. These histone readers provide scaffolds to attract components of the transcriptional machinery to histone acetylation marks. Pharmacological inhibition of BET proteins results in repression of downstream gene expression, thereby modulating various physiological conditions. Indeed, prototype BET inhibitors such as JQ1 or I-BET attenuated various types of cancer and tissue fibrosis (5–9).

Considering the potential anti-fibrotic properties of BET inhibition, we investigated whether BET inhibitor JQ1 can modulate fibrogenesis in an animal model of SSc, as well as in dermal fibroblasts isolated from patients with diffuse cutaneous SSc (dcSSc). We posit that BET inhibition impedes the expression of pro-fibrotic genes and blocks myofibroblast differentiation in SSc, thereby improves fibrosis. Through whole transcriptomic analysis we revealed a novel mechanism by which BET inhibition exerts its anti-fibrotic effect in SSc. Further, we demonstrated the involvement of BRD4 in mediating the pro-fibrotic effect of BETs and identified BRD4 as a novel therapeutic target for this disease. The novelty of our study is that we performed a comprehensive analysis by employing both functional assays and transcriptomic analysis that allowed us to pinpoint a novel mechanism where BET inhibition affects intracellular calcium and related signaling in SSc fibroblasts. We also showed the anti-fibrotic properties of two novel BRD4 inhibitors, one (AZD5153) currently evaluated in a clinical trial (NCT03205176). The effect of these inhibitors has not been examined in other fibrotic models before.

## Results

### The anti-fibrotic effects of JQ1 in bleomycin-treated mice

To investigate the anti-fibrotic effect of BET inhibition, we used a pan-BET inhibitor, JQ1, in an animal model of fibrosis. We observed approximately 3.5% weight reduction in the bleomycin/vehicle group and a 11.4% weight reduction in the bleomycin/JQ1 group. All animals were active without apparent distress throughout the course of the experiment. Daily injections of bleomycin in a defined area in the back of mice increased dermal thickness and collagen accumulation (**Figure 1A**). Daily JQ1 oral gavage prevented skin fibrosis in bleomycin-treated mice, as significantly attenuation of dermal thickness and collagen was observed. In addition, immunofluorescent staining revealed increased αSMA- and F4/80-positive cells in bleomycin-treated mice, with F4/80 staining reaching statistical significance (**Figure 1B**). In addition, JQ1 treatment reduced αSMA- and F4/80-positive cells, with αSMA staining reaching statistical significance. These results were also reflected at the mRNA level, as pro-fibrotic genes including *Acta2* and *Col1a1* that were significantly elevated in bleomycin-treated mice, were downregulated in the presence of JQ1 (**Figure 1C**). Bleomycin also induced significant increase in *Il6* and *Ccl2*, and both were significantly downregulated with JQ1 treatment. In contrast, *Ctgf* and *Cxcl10* were not significantly altered by bleomycin or JQ1.

**Figure 1.**
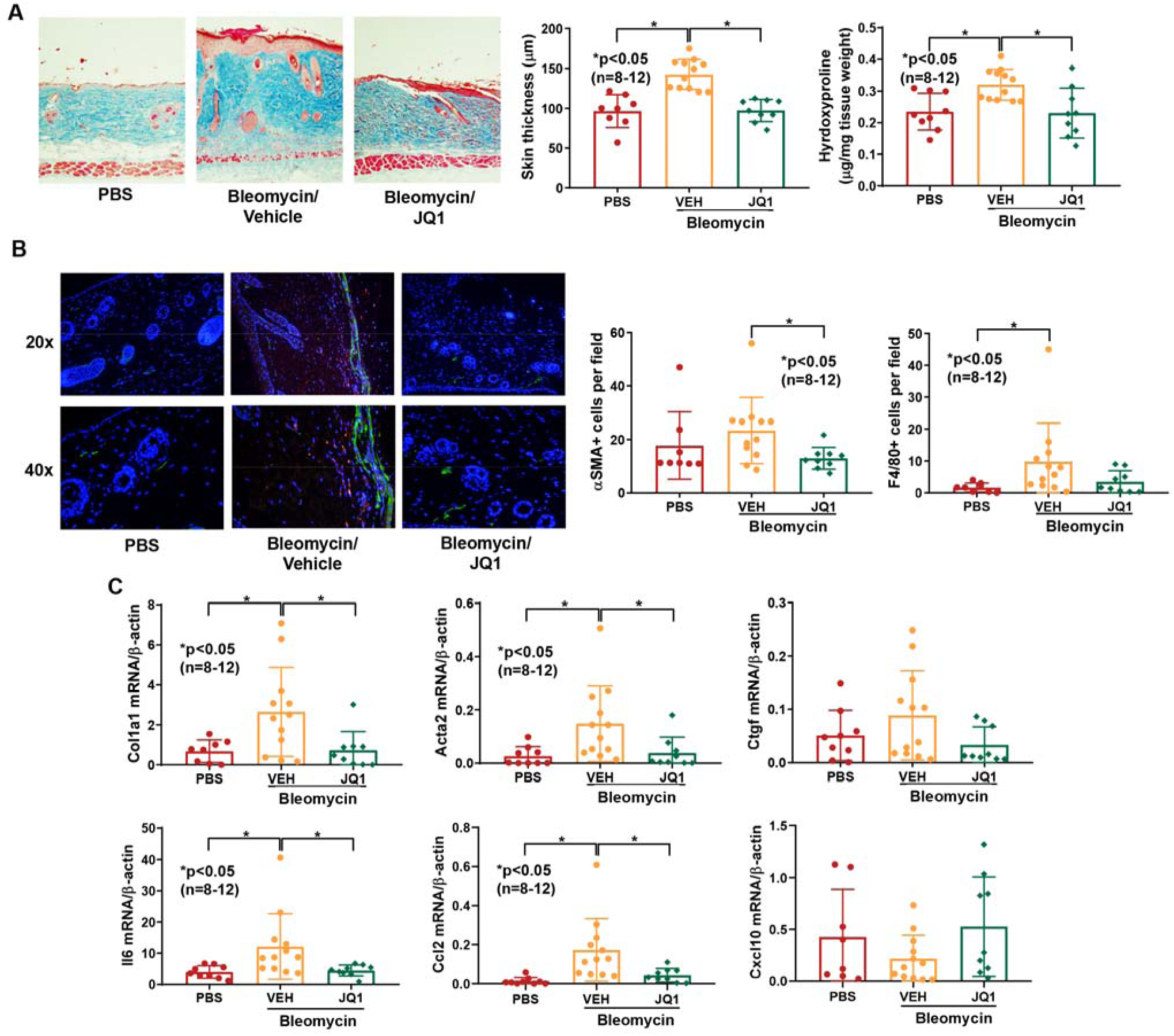
BET inhibition by JQ1 prevents bleomycin-induced skin fibrosis in mice. (**A**) Bleomycin-treated mice showed increased dermal thickness and hydroxyproline content in skin, and JQ1 (50mg/kg) efficiently prevented skin fibrosis in these mice. Original magnification: x100; (**B**) Immunofluorescent staining of skin sections showed that JQ1 significantly decreased αSMA-positive cells (green) in bleomycin-treated mice compared to the vehicle (VEH) group. Similar results were found quantifying F4/80-positive cells (red). Nuclei were stained with DAPI (blue). Original magnification: x200-400; (**C**) *Acta2, Col1a1*, *Il6*, and *Ccl2* were significantly elevated in bleomycin-treated mice, while significantly reduced when JQ1 was given. n=number of animals (PBS: n=8, VEH: n=12, JQ1: n=9). Results are expressed as mean +/- SD and p<0.05 was considered significant. Significance was determined by one-way ANOVA (**A**, **C**) and Kruskal-Wallis test (**B**, **C**).

### Anti-fibrotic properties of BET inhibition in dermal fibroblasts

To further evaluate the effect of BET proteins in SSc fibrosis, we treated dcSSc fibroblasts with JQ1 at various doses (**Supplemental Figure 1**). Given that the highest dose (22 µM) had minimal effect on *CTGF* while upregulated *TGFB1*, suggesting a potential off-target effect, we found that 1 μM appears to be the optimal dose to use. We evaluated the expression of pro-fibrotic genes *COL1A1*, *ACTA2, CTGF,* and *TGFB1,* as well as collagen-degrading enzyme *MMP1* and its inhibitor *TIMP3.* Inhibition of BET by JQ1 led to a dose-dependent decrease in fibrotic markers including *COL1A1*, *ACTA2, and CTGF* (**Supplemental Figure 1** and **Figure 2A**). In addition, JQ1 also increased *MMP1*, which is critical for collagen turnover, while it had minimal effect on *TIMP3*. Interestingly, JQ1 significantly increased *TGFB1*. This might be an off-target effect of the inhibitor at high doses.

**Figure 2.**
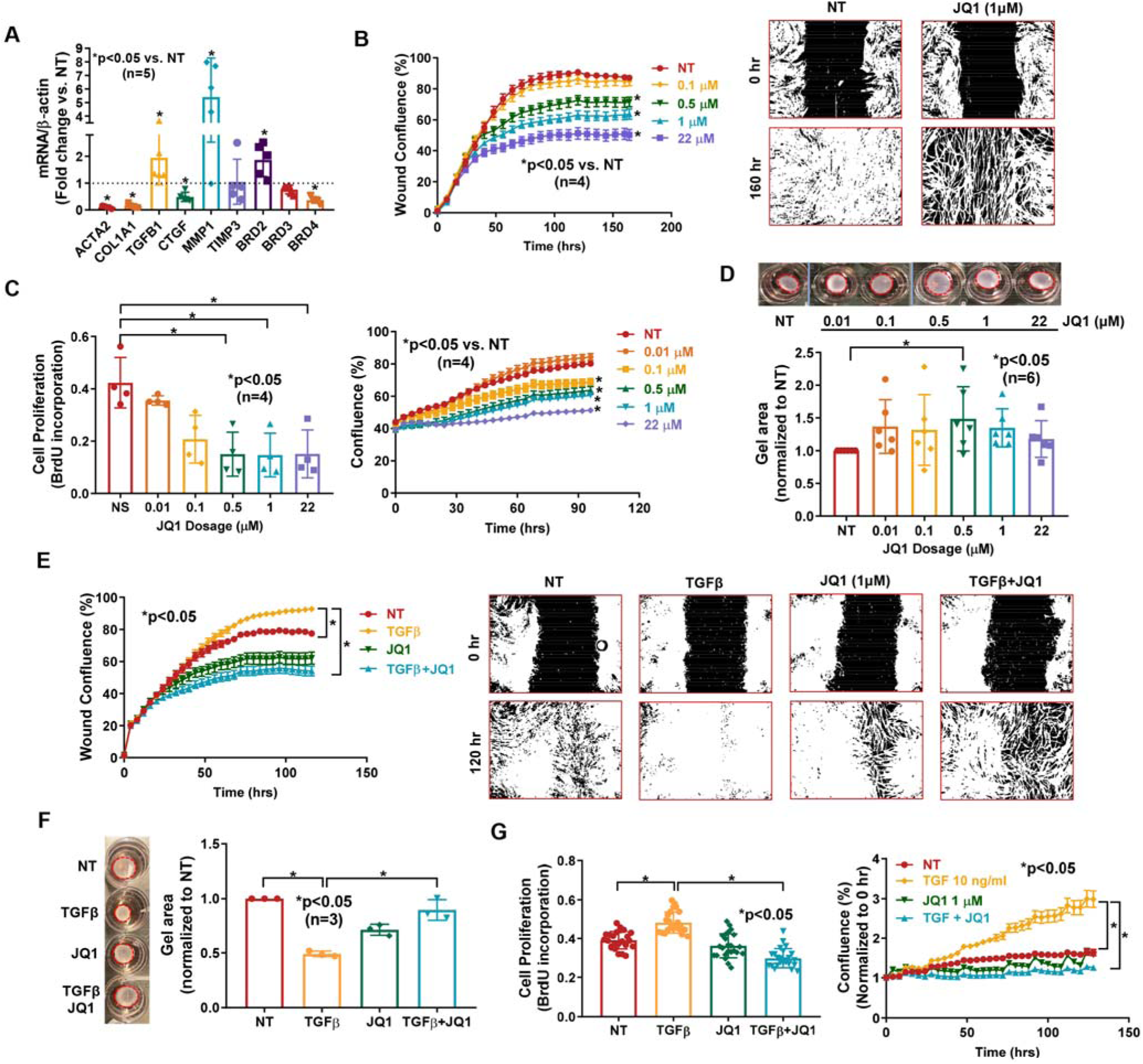
Inhibition of BETs shows potent anti-fibrotic properties in dcSSc fibroblasts. (**A**) At 1 µM, JQ1 significantly downregulated *ACTA2*, *COL1A1*, *CTGF* and *BRD4* in dcSSc fibroblasts, and upregulated *MMP1*, *TGFB1*, and *BRD2*. JQ1 did not affect *TIMP1* and *BRD3* expression. n=5 patients; (**B**) Migration of dcSSc fibroblasts was significantly inhibited by JQ1. Wound confluence indicates the area occupied by cells migrated into the wound gap. Representative pictures of JQ1 at 1 µM was shown. n=4 patients; (**C**) Inhibition of BETs by JQ1 significantly reduced cell proliferation of dcSSc fibroblasts. Cell growth was analyzed by BrdU uptake in cells or monitored by IncuCyte® Live-cell imaging. n=4 patients; (**D**) Gel contraction by dcSSc fibroblasts was inhibited by JQ1 at 0.5 µM. n=6 patients; (**E-G**) Cell migration (n=4), proliferation (n=4), and gel contraction (n=3) significantly increased after TGFβ treatment in normal dermal fibroblasts, and can be inhibited by co-incubation of 1 µM of JQ1. n=number of patients. Results are expressed as mean +/- SD or mean +/- SEM (time-courses in **B**, **C**, **E**, **F**) and p<0.05 was considered significant. Significance was determined by unpaired t test/Mann-Whitney test (**A**), two-way ANOVA (**B**, **C**, **E**, **G**), one-way ANOVA (**F**), and Kruskal-Wallis test (**D, G**)

To further characterize the anti-fibrotic effect of BET inhibition in dcSSc fibroblasts, we performed several functional assays. Treatment with JQ1 dose-dependently reduced migration and proliferation of dcSSc fibroblasts (**Figure 2B** and **2C**). The mechanism of reduction of cell proliferation by JQ1 might be in part due to its effect on inducing apoptosis, as JQ1 dose dependently increased apoptotic cells as indicated by green fluorescence released by activated caspase-3/7 using IncuCyte® (**Supplemental Figure 2A**). However, this effect appears to be delayed, as apoptotic cells appeared after 40 hours while differences in cell proliferation became evident after 30 hours of JQ1 treatment (**Figure 3C** **right panel** vs. **Supplemental Figure 2A**). This suggests that other mechanisms are involved in the effect of JQ1 on cell growth. To further analyze the effect of JQ1 on cell growth, we used Vybrant DyeCycle Violet stain to analyze cell cycle distribution in dcSSc fibroblasts. JQ1 induced a reduction in cells in the S-phase and accumulation of cells in the G1/G0 phase (**Supplemental Figure 2B**). Treatment of dcSSc fibroblasts with JQ1 also resulted in decrease in gel contraction (**Figure 2D**), with 0.5 µM being the most effective concentration.

**Figure 3.**
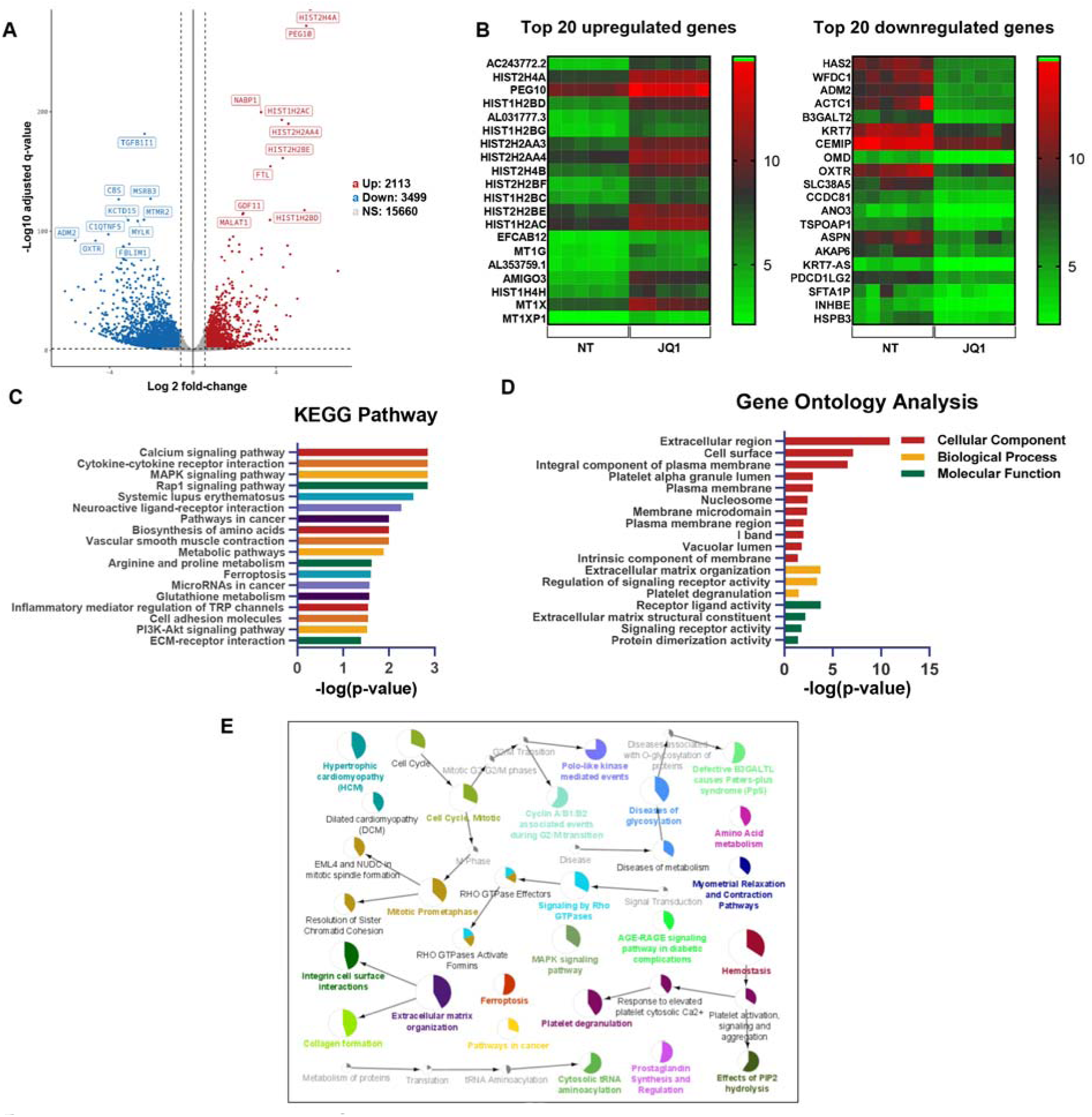
JQ1-treatment followed by RNA-seq identified targets important for ECM remodeling, cell cycle regulation, and signaling in dcSSc fibroblasts. (**A**) A total of 6 patient pairs were used for RNA-seq analysis. In total, 5612 genes were significantly differentially expressed by JQ1 treatment in dcSSc fibroblasts (3499 downregulated and 2113 upregulated). (**B**) Top 20 upregulated and downregulated genes were shown. (**C**) The 18 most significantly enriched KEGG pathways after treatment with JQ1 in dcSSc fibroblasts are shown. (**D**) Gene ontology analysis of the differentially expressed genes after JQ1 treatment in dcSSc fibroblasts. (**E**) Network of functional categories represented by JQ1-associated changes in dcSSc fibroblasts based on functional analysis. The node size corresponds with the level of significance (p-values range from <0.05 to <0.0005) of the term it represents within the network. Colored portions of the pie charts represent the number of genes identified from the RNA-seq experiment that are associated with the term.

Since TGFβ is critical in promoting fibrosis in SSc, we treated normal dermal fibroblasts with TGFβ to induce a myofibroblast phenotype. This was confirmed with an increase in cell migration, proliferation, as well as gel contraction in these TGFβ-treated normal fibroblasts compared to non-treated controls (**Figure 2E-G**). Co-treatment with JQ1 significantly reduced TGFβ-induced migration, proliferation, and gel contraction in normal dermal fibroblasts.

### Transcriptome analysis

To further explore the mechanisms involved in BET inhibition in SSc fibrosis, mRNA-seq was performed on dcSSc fibroblasts in the absence or presence of 1μM JQ1. This analysis identified 2113 upregulated and 3499 downregulated genes affected by JQ1 (**Figure 3A** and **Supplemental Figure 3**). The excess number of downregulated genes with BET inhibition indicates that BET proteins primarily act as transcription activators in SSc fibroblasts, echoing the findings in other tissues indicating that histone readers are indeed transcription activators. The top 20 upregulated and downregulated genes are shown in **Figure 3B**. In addition, RNA-seq results confirmed expression changes observed using qPCR in fibrosis-related genes (**Figure 2A**), where *ACTA2* and *COL1A1* were significantly downregulated while *TGFB1* and *BRD2* were significantly upregulated at 1μM of JQ1 treatment (**Supplemental Figure 4**). Further pathway enrichment analysis of the differentially expressed genes regulated by JQ1 revealed that these genes are enriched in KEGG pathways including calcium signaling pathway, cytokine-cytokine receptor pathway, MAPK and Rap1 signaling pathways, as well as metabolic pathways, among others (**Figure 3C** and **Supplemental Table 1**). In addition, gene ontology analysis showed that BET proteins are involved in a wide spectrum of cellular components, biological processes, and molecular functions known to play critical roles in fibroblast activation (**Figure 3D**). These include ECM, plasma membrane, and signaling receptor activity. To better visualize the network of functional categories represented by JQ1-associated changes, we performed additional functional enrichment analysis using ClueGO/CluePedia by incorporating the KEGG, REACTOME, and WikiPathways. As shown in **Figure 3E**, networks related to cell cycle regulation, ECM organization, signaling by Rho GTPases, diseases of glycosylation, ferroptosis, and homeostasis, among others, are significantly enriched. We also examined the subcellular localization of the enriched pathways using the cerebral layout tool implemented in Cytoscape. The subcellular localization of JQ1-affected pathways was skewed toward extracellular and plasma membrane (**Supplemental Figure 5**). This is not surprising, as functions related to ECM and homeostasis, as well as signaling pathways, were among the most enriched (as demonstrated by the size of the nodes). Indeed, when the differentially expressed genes were overlaid on the KEGG: 04512 ECM-receptor interaction pathway, many ECM genes and their binding partners were affected by JQ1, with majority of these genes downregulated (**Supplemental Figure 6**). In the intracellular compartment and the nucleus, cell cycle regulation is the main enriched pathway (**Supplemental Figure 5**). These results echo our functional data shown in **Figure 2** and **Supplemental Figure 2**, as JQ1 showed strong anti-proliferative and anti-fibrotic effects in these cells. Finally, as expected, the predicted upstream regulator analysis showed JQ1 as the most significant upstream chemical, as 1521 genes from our transcriptomic analysis overlapped with reported JQ1-target genes (**Supplemental Figure 7**). This not only validates the JQ1 inhibition condition used and sequencing analysis employed in this study, but also shows that genes affected by BET inhibition are common among different biological systems and cell types.

### BET inhibition affects the calcium signaling pathways in dcSSc fibroblasts

As shown in **Figure 3C**, the most significant enrichment of the differentially expressed genes in the KEGG pathways was the calcium signaling pathway. To better visualize the location of the differentially expressed genes, we overlaid them in the pathway. JQ1 appears to affect genes involved in calcium (Ca^2+^) channels on the plasma membrane, Ca^2+^ receptors that are located in the endoplasmic/sarcoplasmic reticulum, Ca^2+^-related signaling pathways, as well as Ca^2+^-dependent downstream effectors (**Figure 4A**). All of the differentially expressed genes enriched in this pathway were listed in **Figure 4B**. 15 out of the 59 genes were upregulated by JQ1, again pointing BET proteins to be transcriptional activators. As calcium signaling affects fibroblast function and myofibroblast transformation (10), we examined whether JQ1 affects intracellular Ca^2+^ concentration in dcSSc fibroblasts. As shown in **Figure 4C**, JQ1 significantly reduced intracellular Ca^2+^ levels in dcSSc fibroblasts.

**Figure 4.**
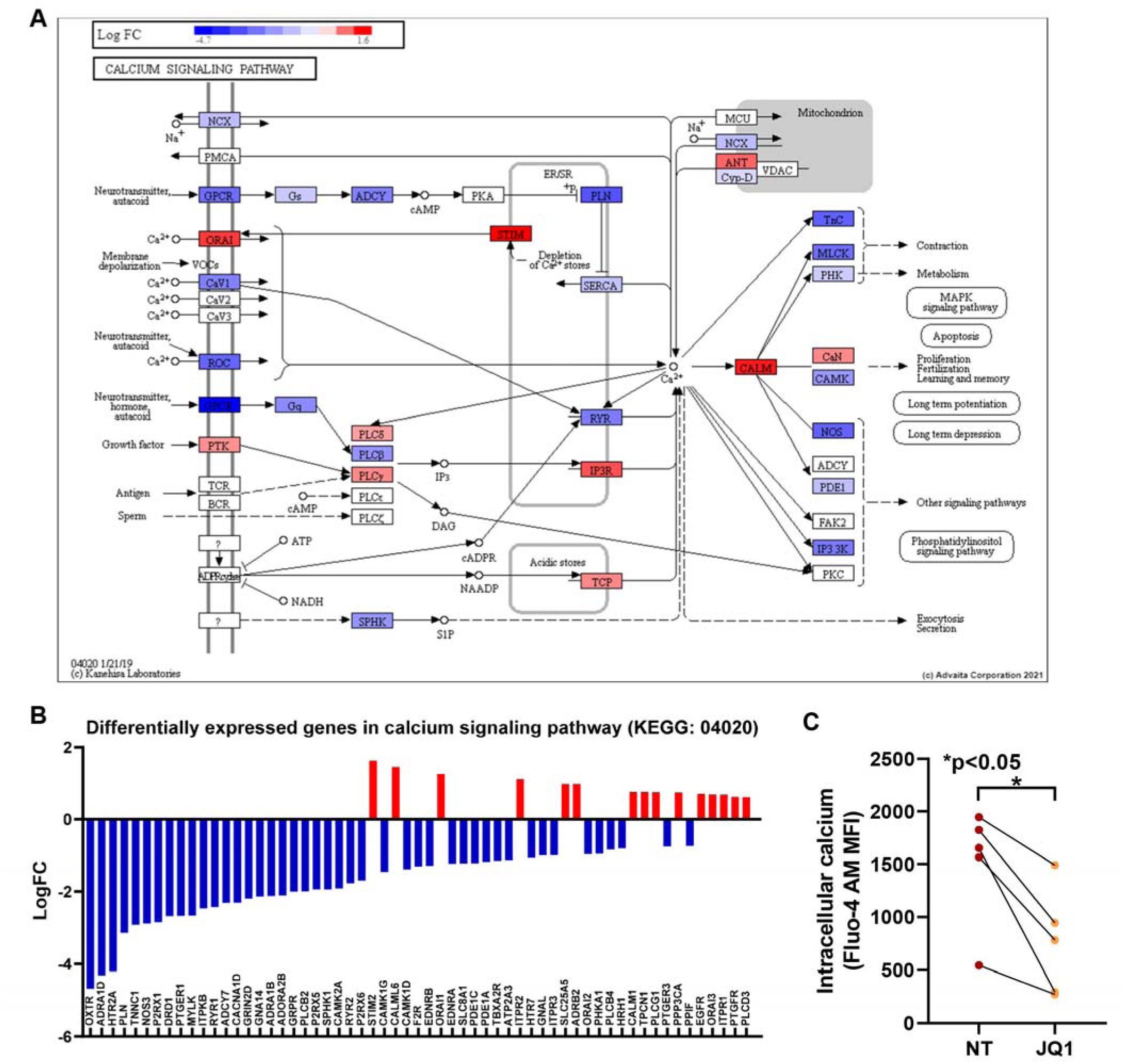
BET inhibition affects calcium-related pathways and intracellular Ca^2+^ in dcSSc fibroblasts. (**A**) The calcium signaling pathway (KEGG: 04020) diagram is overlaid with the expression changes of each gene. The legend describes the values on the gradient. Downregulation is shown in blue, while upregulation is in red. For legibility for the figure, one gene may be presented at multiple boxes in the diagram. In addition,one box may represent multiple genes in the same gene family. For each gene family, the color corresponding to the gene with the highest absolute fold change is displayed. For example, the ORAI box depicts significant upregulation of *ORAI1* and *ORAI3* and downregulation of *ORAI2*. It is shown in red, since *ORAI1* shows the most significant fold change among the three. (**B**) All the differentially expressed genes in calcium signaling pathway (KEGG: 04020) are ranked based on their absolute value of log fold change. Upregulated genes are shown in red, downregulated genes are shown in blue. n=6 patient pairs; (**C**) BET inhibition by JQ1 significantly reduced intracellular calcium levels in dcSSc fibroblasts. n= 5 patients. Results are expressed as mean +/- SD and p<0.05 was considered significant. Significance was determined by paired t test (**C**).

### JQ1 mediates its anti-fibrotic effect in part by affecting calcium signaling in dcSSc fibroblasts

To further examine the involvement of intracellular Ca^2+^ in the anti-fibrotic effect of JQ1, we cultured dcSSc fibroblasts in the presence or absence of Ca^2+^, and treated the cells with or without JQ1. Comparing the non-treated groups in the presence or absence of Ca^2+^ in culture media, the expression of *ACTA2* and *COL1A1*was significantly lower in cells cultured in media without Ca^2+^ (**Figure 5A**). JQ1 significantly downregulated *ACTA2* and *COL1A1* in Ca^2+^-containing media, while its effect on *COL1A1* was diminished in media without Ca^2+^. Using BAPTA-AM, an intracellular Ca^2+^-chelating reagent, we also showed that depleting Ca^2+^ in cells decreased *ACTA2* and *COL1A1* expression, and the inhibitory effect of JQ1 on these genes was blocked (**Figure 5B**). To further demonstrate the involvement of Ca^2+^ in the anti-fibrotic effect of JQ1, we focused on *CAMK2A*, which encodes Ca^2+^/calmodulin-dependent protein kinase II α (CaMKII-α), in subsequent studies. This gene is central for Ca^2+^-mediated effects in cells and was significantly downregulated by JQ1 (**Figure 4A-B**). We further confirmed the effect of JQ1 on *CAMK2A expression* by qPCR (**Figure 5C**). Overexpression of *CAMK2A* in dcSSc fibroblasts blocked the inhibitory effect of JQ1 on *COL1A1* expression (**Figure 5C**) but not on *ACTA2*. Since CaMKII is involved in cell cycle (11), we examined whether *CAMK2A* is involved in JQ1-mediated anti-proliferative effect in dcSSc fibroblasts. Indeed, we showed that *CAMK2A* appears to be involved, at least in part, in the anti-proliferative effect of JQ1, as overexpression of *CAMK2A* resulted in approximately 25% reduction in cell growth by JQ1 compared to 50% in the control group (**Figure 5D**). Notably, *CAMK2A* overexpression in dcSSc fibroblasts not only resulted in elevated levels of *ACTA2* and *COL1A1* but also increased cell proliferation, suggesting that this gene plays a critical and non-redundant role in promoting SSc fibrosis. These results support the involvement of Ca^2+^ and Ca^2+^-related pathways in the anti-fibrotic effect of JQ1, specifically on collagen expression and cell proliferation.**BET expression in SSc.** We examined the expression of BET proteins in dermal fibroblasts. Out of the four BETs, BRD-t is expressed predominantly in the testis and therefore excluded from our study. At the mRNA level, only *BRD4* was significantly upregulated in dcSSc fibroblasts compared to normal fibroblasts (**Supplemental Figure 8A**). At the protein level, however, the expression of the BETs was variable in dcSSc fibroblasts but no difference between normal and dcSSc was seen (**Supplemental Figure 8B**).

**Figure 5.**
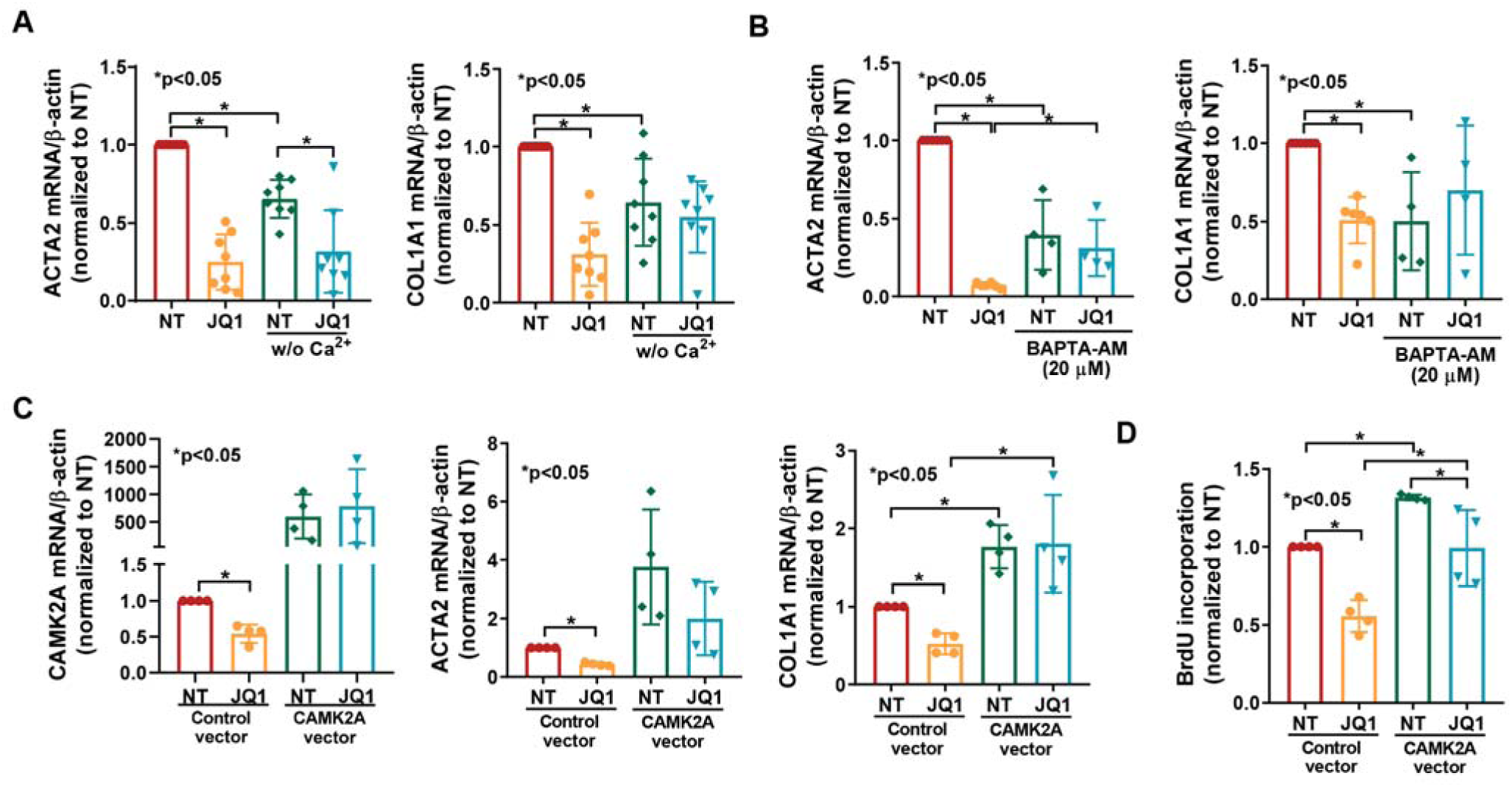
The anti-fibrotic effect of JQ1 is abolished by the sequestration of intracellular Ca^2+^ and overexpression of *CAMK2A* in dcSSc fibroblasts. (**A**) Cells cultured in Ca^2+^-free media showed lower levels of *ACTA2* and *COL1A1* compared to ones cultured in Ca^2+^-containing media. The effect of JQ1 on COL1A1 is blocked in cells cultured without Ca^2+^. n=8 patients; (**B**) BAPTA-AM, an intracellular Ca^2+^ chelating reagent, significantly decreased *ACTA2* and *COL1A1* expression in dcSSc fibroblasts. It also blocked the effect of JQ1. n=4 patients; (**C**) JQ1 downregulated *CAMK2A* expression in dcSSc fibroblasts. Overexpression of *CAMK2A* not only increased *ACTA2* and *COL1A1* expression, it also blocked the anti-fibrotic effects of JQ1. n=4 patients; (**D**) *CAMK2A* is critical for cell proliferation, as overexpression of this gene enhanced cell proliferation. The inhibitory effect of JQ1 on cell proliferation was blocked by *CAMK2A* overexpression. n=4 patients. Results are expressed as mean +/- SD and p<0.05 was considered significant. Significance was determined by one-way ANOVA.

### BRD4 is pro-fibrotic

Since JQ1 is a pan-BET inhibitor, we next performed a set of experiments to determine which BET is responsible for the anti-fibrotic effect of JQ1. JQ1 significantly induced *BRD2* and decreased *BRD4*, while it had minimal effect on *BRD3* (**Supplemental Figure 1** and **Figure 2A**). We then knocked down BRD2, BRD3, or BRD4 individually and examined the expression of genes involved in fibrosis. As shown in **Figure 6A**, knockdown of BRD2 resulted in significant elevation of *COL1A1*, while knockdown of BRD4 led to significant reduction of *COL1A1*, *ACTA2*, *TGFB1*, *CTGF*, and *MMP1*. BRD3 knockdown decreased *TIMP3* while it had no effect on the other genes examined. In addition, BRD4 knockdown led to significant reduction in gel contraction (**Figure 6B**), suggesting the BRD4 is pro-fibrotic, and that JQ1, by significantly reducing BRD4 expression in dcSSc fibroblasts, inhibits fibrosis in SSc.

**Figure 6.**
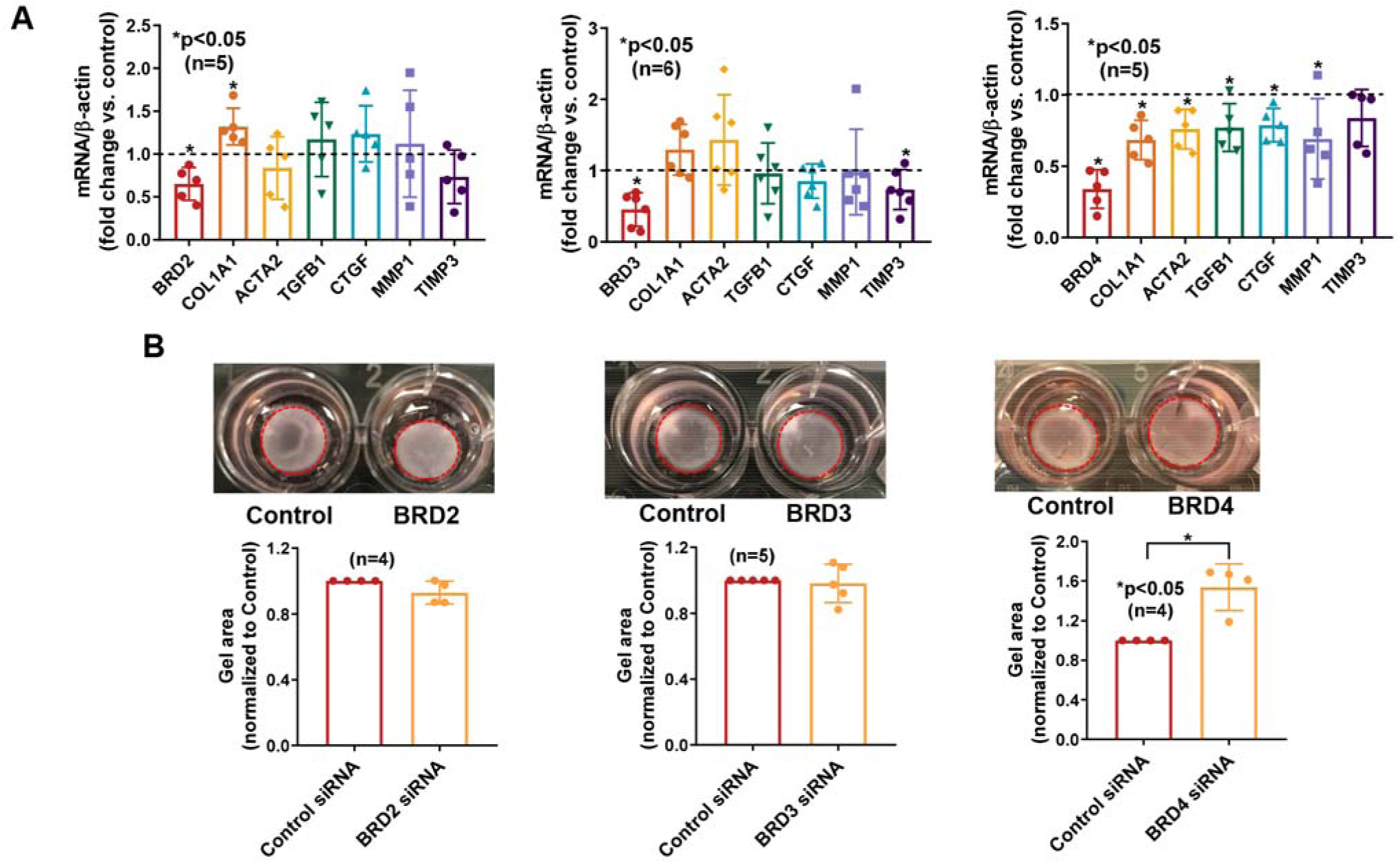
BRD4 mediates the pro-fibrotic effect of BETs in dcSSc fibroblasts. (**A**) BRD2 knockdown in dcSSc fibroblasts led to upregulation of *COL1A1* while BRD3 knockdown resulted in downregulation of *TIMP3*. Knockdown of BRD4 resulted in downregulation of *COL1A1*, *ACTA2*, *TGFB*, *CTGF*, and *MMP1*. n=5-6 patients; (**B**) BRD4 knockdown significantly inhibited gel contraction in dcSSc fibroblasts while BRD2 or BRD3 knockdown had minimal effect on gel contraction. n=4-5 patients. Results are expressed as mean +/- SD and p<0.05 was considered significant. Significance was determined by unpaired t test or Mann-Whitney test (**A, B**).

To further confirm the involvement of BRD2 and BRD4 in myofibroblast function, we treated dcSSc fibroblasts with specific BRD2 inhibitor (B1C1) or BRD4 inhibitors (AZD5153 or ARV825). As shown in **Figure 7A**, inhibition of BRD2 by BIC1 did not affect dcSSc fibroblast proliferation, while BRD4 inhibition by either AZD5153 orARV825 significantly inhibited proliferation at both 1 and 10 µM. In addition, dcSSc fibroblasts treated with BRD4 inhibitors ARV825 or AZD5153 significantly reduced *ACTA2* and *COL1A1* while BRD2 inhibitor BIC1 did not have any effect (**Figure 7B**). Similar results were observed at the protein level (**Figure 7C**). To further determine the anti-fibrotic effect of BRD4 inhibition in vivo, we induced skin fibrosis by bleomycin in mice and dosed the animals with either ARV825 or AZD5153. We found that both BRD4 inhibitors prevented bleomycin-induced skin fibrosis in mice, shown by dermal thickness, αSMA-positive cells, and hydroxyproline content (**Figure 7D**). These results further confirm the involvement of BRD4 in myofibroblast transformation in dcSSc fibroblasts and in the bleomycin mouse model.

**Figure 7.**
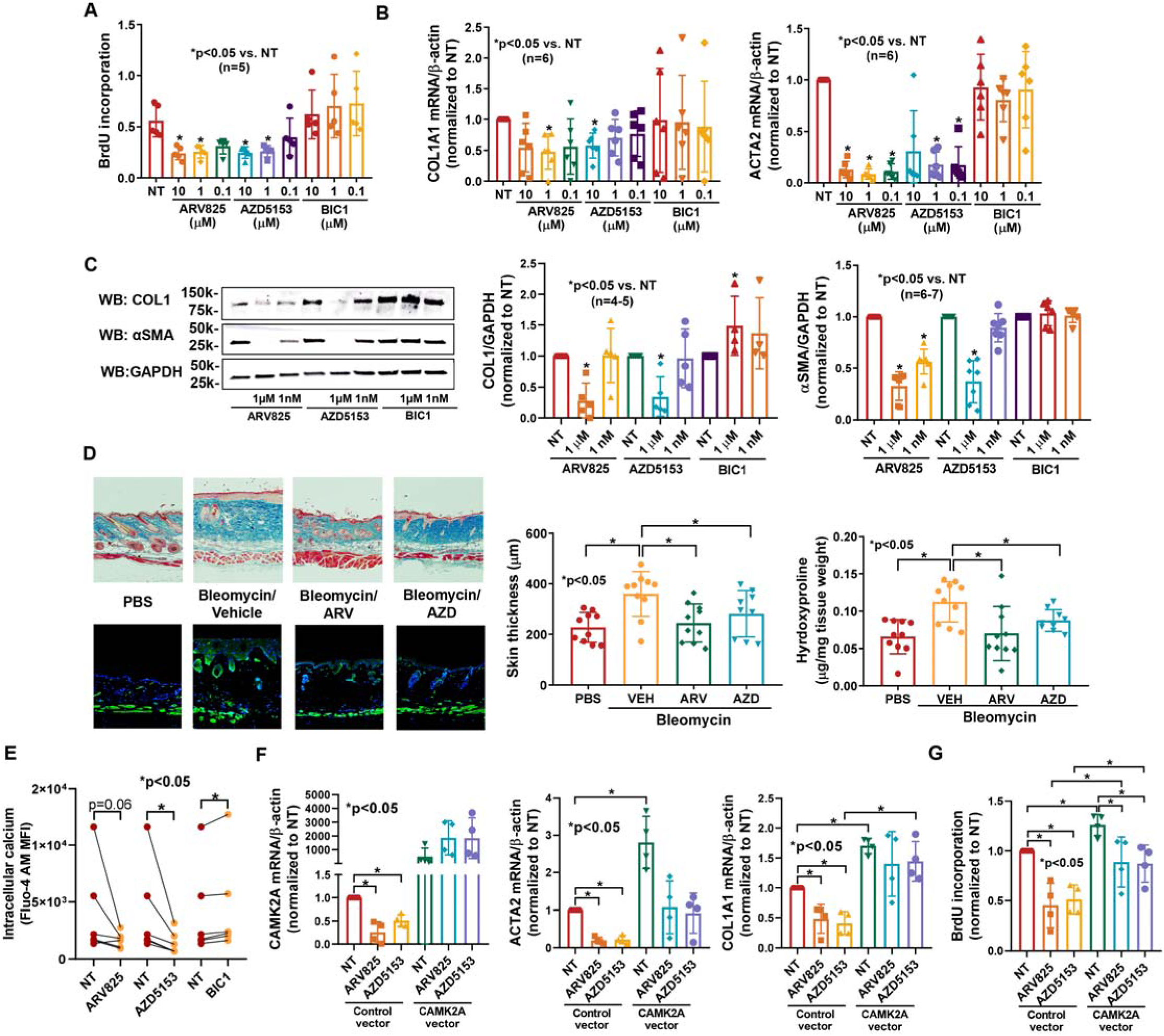
BRD4 inhibitors show prominent anti-fibrotic effects in vitro and in vivo. (**A**) Treating dcSSc fibroblasts with specific BRD2 inhibitor BIC1 had minimal effect on cell proliferation, while inhibition of BRD4 using BRD4 inhibitors AZD5153 or ARV825 significantly reduced cell proliferation at concentrations of 1 and 10 µM. n=5 patients; (**B**) Inhibition of BRD4 by ARV825 or AZD5153 in dcSSc fibroblasts significantly reduced both *ACTA2* and *COL1A1* expression, while blockade of BRD2 by BIC1 had no effect.n=6 patients; (**C**) BRD4 inhibitors significantly decreased αSMA (n=7) and COL1 (n=5) in dcSSc fibroblasts while BRD2 inhibition by BIC1 had minimal effect on αSMA (n=6) but increased COL1 at 1 μM (n=4, n=number of patients); (**D**) Bleomycin-treated mice showed increased dermal thickness and hydroxyproline content in skin, and ARV825 or AZD5153 efficiently prevented skin fibrosis in these mice. Immunofluorescent staining of αSMA-positive cells (green) was shown. Nuclei were stained with DAPI (blue). Original magnification: x40-100. n=9-10 mice; (**E**) BRD4 inhibitor AZD5153 significantly decreases intracellular Ca^2+^ while BRD2 inhibitor BIC1 significantly increased it. n=6 patients; (**F**) Overexpression of *CAMK2A* resulted in significant increase in *ACTA2* and *COL1A1* expression while blocked the effect of AZD5153 on *COL1A1* expression. n=4 patients; (**G**) *CAMK2A* overexpression significantly increased cell proliferation while it abolished the effect of BRD4 inhibition on cell proliferation. n=4 patients. Results are expressed as mean +/- SD and p<0.05 was considered significant. Significance was determined by one-way ANOVA (**A**, **B**, **C, D, F, G**), Kruskal-Wallis test (**A**, **B**, **C**), and Wilcoxon test (**E**).

### BRD4 mediates its fibrotic properties through Ca^2+^ signaling in dcSSc fibroblasts

To further investigate whether BRD4 inhibitors affect intracellular Ca^2+^ in dcSSc fibroblasts similar to what was observed with JQ1, we measured intracellular Ca^2+^ levels in the presence or absence of the BRD4 inhibitors. We found that both inhibitors decreased intracellular Ca^2+^ levels after 48 hours of treatment, with AZD5153 reaching statistical significance (**Figure 7E**). Interestingly the BRD2 inhibitor BIC1 significantly increased intracellular Ca^2+^, suggesting that this BET isoform could act in an opposite way compared to BRD4 in dcSSc fibroblasts. We further showed that both BRD4 inhibitors downregulated *CAMK2A* in dcSSc fibroblasts (**Figure 7F**). In addition, the anti-fibrotic effects of both AZD5153 and ARV825 in dcSSc fibroblasts were dependent on *CAMK2A*. Overexpression of *CAMK2A* in these cells appeared to block the inhibitory effect of ARV825 and AZD5153 on fibrotic gene expression, with *COL1A1* reaching statistical significance for AZD5153 (**Figure 7F**). BRD4 inhibitors also blocked cell proliferation by 50% in control cells but only by approximately 25% in the *CAMK2A* overexpressing cells, suggesting that *CAMK2A* is involved in BRD4-mediated effects on cell growth (**Figure 7G**).

## Discussion

The increased understanding of the effect of epigenetic aberrations on gene transcription has led the field to a better appreciation of the role of transcriptional dysregulation in initiating and perhaps maintaining SSc fibrosis. In this study we performed an in-depth examination of BET inhibition in SSc fibrosis. Our in vivo studies showed that administration of JQ1 prevented skin fibrosis induced by bleomycin in mice. We further demonstrated that BET inhibition by JQ1 suppressed expression of many pro-fibotic genes in dcSSc fibroblasts and reversed the established progression of myofibroblast differentiation in vitro. Transcriptomic analysis of JQ1-treated cells not only showed that differentially expressed genes are enriched in ECM- and cell cycle-related pathways but also revealed the involvement of calcium signaling pathway as a novel anti-fibrotic mechanism. Moreover, results from siRNA knockdown experiments confirmed the role of BRD4 in SSc fibrosis. This was further confirmed using BRD4 specific inhibitors. Functional studies showed that Ca^2+^ and Ca^2+^-related gene *CAMK2A* plays a critical role in the anti-fibrotic effects of JQ1, AZD5153, and ARV825. Together, these results suggest that BRD4 is critically involved in promoting fibrosis in SSc, and inhibition of BRD4, and perhaps targeting intracellular Ca^2+^, would be effective treatments for this disease.

JQ1, a first in-class BET inhibitor, competitively binds to the acetyl-lysine recognition areas of these proteins, and displaces them from acetylated chromatin, thereby repressing transcription of target genes. Early studies suggested that JQ1 exerted potent anti-proliferative properties in multiple myeloma via cell-growth arrest and senescence in a c-MYC-dependent manner (12). Since then, more studies have shown the benefits of JQ1 in other proliferative disorders (5–8). In addition, this pan-BET inhibitor has demonstrated great efficacy in blocking fibrotic progression in a range of fibrosis models (5–8). In a lung fibrosis model, JQ1 significantly reduced collagen deposition in bleomycin-treated mice compared to control mice (8). This drug also ameliorated the phenotypic changes of lung fibroblasts from idiopathic pulmonary fibrosis patients. We showed in this study that JQ1 inhibits proliferation of dcSSc dermal fibroblasts through inducing cell cycle arrest and apoptosis. It also exhibits potent anti-fibrotic effects in patient-derived cells and in the bleomycin skin fibrosis model. The functional changes in cell proliferation are also confirmed from the RNA-seq results, as genes enriched in cell cycle, mitosis, and ferroptosis pathways are highlighted in the pathway analysis (**Figure 3E**). Our results are consistent with the findings from Shin et al, where they showed that JQ1 repressed collagen expression in SSc skin organ culture, likely due to the increase in MMP1 (13).

BET proteins are located in the nucleus. Because of their involvement in various cellular processes, BETs have been shown to participate in tumor development, autoimmunity, infections, and inflammation (14–17). Specifically, BRD4 has been studied extensively for its role in gene transcription, including regulation (through acetylated histones), initiation (via engaging RNA polymerase II), and elongation (by interaction with P-TEFb)(18). Indeed, BRD4 is reported to control various fibrosis-related genes, including *ACTA2* and *COL1A1*, shown from BRD4 knockdown studies or ChIP analyses (5, 7, 8, 19). This is not surprising, as both genes are reported to possess acetylated histones in their promoter region in fibroblasts (**Supplemental Figure 9**). In dcSSc fibroblasts, inhibition of BETs by JQ1 downregulated both *ACTA2*and *COL1A1*, and this appears to be mediated through blockade of BRD4, as BRD4 knockdown or inhibition led to decreased *ACTA2* and *COL1A1* levels, while knockdown of BRD2 or BRD3 did not. Interestingly, *TGFB1*, which was significantly upregulated by JQ1, was downregulated in BRD4 knocked down cells. Similar results were seen in *MMP1* expression. This discrepancy suggests that JQ1, as a pan-BET inhibitor, has off-target effects compared to BRD4 knockdown, especially at the higher concentrations. This could potentially explain the U-shaped dose-response curve of JQ1 in the gel contraction assay. Of note, our results showing that BRD2 knockdown leads to *COL1A1* upregulation are consistent with previous reports (19). Since JQ1 dose-dependently increased *BRD2* expression in dcSSc fibroblasts, it is possible that the anti-fibrotic effect of JQ1 is partly mediated by enhancing the anti-fibrotic potential of BRD2. However, BRD2 inhibition, although trending to induction of fibroblast migration and pro-fibrotic gene expression (**Figure 7A-C**), had no significant effect on enhancing myofibroblast functions. Furthermore, BIC1 significantly increased intracellular Ca^2+^ levels in dcSSc fibroblasts, in contrast to BRD4 inhibition.

To gain mechanistic insight into the anti-fibrotic effect of BET inhibition in SSc, we performed mRNA-seq in JQ1-treated dcSSc fibroblasts. In addition to pathways in cell cycle regulation, ECM, and intracellular signaling, our analysis also revealed many novel pathways, including calcium signaling, metabolism, ferroptosis, and prostaglandin synthesis and regulation (**Figure 3E**). As these pathways have been implicated in various fibrotic conditions, it is possible that they are involved in the anti-fibrotic effect of BET inhibition in dcSSc fibroblasts. Specifically, the calcium signaling pathway, which is the most enriched pathway in our analysis (**Figure 3C**), has been shown to be critical in various fibrotic conditions (10, 20, 21). Intracellular Ca^2+^ levels are controlled by Ca^2+^-permeable channels in the plasma membrane (voltage-operated channels and receptor-operated channels), Ca^2+^ release from the endoplasmic/sarcoplasmic reticulum, as well as Ca^2+^ extrusion pumps. The increase in intracellular Ca^2+^ levels can activate various pathways that are involved in many physiological functions (22–25). Studies show that interference of intracellular Ca^2+^ levels through blockade of Ca^2+^ channels or receptors has potential anti-fibrotic effects in various models (26–29). In dcSSc fibroblasts, JQ1 treatment significantly reduced Ca^2+^ levels in these cells (**Figure 4C**). This is possibly mediated by significant downregulation of calcium channels and receptors on the plasma membrane, including *P2RX1* and *P2RX5* (encoding for purinergic receptor P2X1 and 5), *CACNA1D* (calcium voltage-gated channel subunit alpha1D), and *TPCN1* (voltage-dependent calcium channel protein TPC1), as well as ones on the endoplasmic/sarcoplasmic reticulum, such as *RYR1* (Ryanodine receptor 1) (**Figure 4B**). Interestingly, certain Ca^2+^ channels, Ca^2+^ extrusion pumps, and Ca^2+^-release transporters that increase intracellular Ca^2+^ were affected by JQ1. It is possible that these events represent a compensatory mechanism that attempt to antagonize the decreased Ca^2+^ levels in these fibroblasts. The identification of Ca^2+^ pumps/carriers/ channels involved in Ca^2+^ entry, as well as their involvement in mediating the fibrotic effect of BET proteins in SSc requires further investigation.

In this study we identified CaMKII, specifically the α isoform, as a critical mediator for the fibrotic effect of BETs and BRD4. CaMKII, comprised of 4 isoforms, is a downstream messenger of the Ca^2+^ signaling pathway. CaMKII activation augmented collagen production in cardiac fibroblasts, while inhibition of CaMKII blocked proliferation, collagen and profibrotic cytokine production (22, 30, 31). We showed similar findings in dcSSc fibroblasts in this study. In addition to cardiac fibrosis, CaMKII is also involved in pulmonary fibrosis (32), ureteral scar formation (33), and renal fibrosis (34). CaMKII has been shown to be associated with FGFR3/FGF9 in SSc fibroblasts, functioning as a downstream mediator for the pro-fibrotic effect of FGFR3 (25). Interestingly, we found that overexpression of *CAMK2A* alone led to significant increase in fibrotic gene expression and cell proliferation in dcSSc fibroblasts, suggesting that this gene itself could play a critical role in SSc fibrosis.

Although JQ1 has been shown to possess desirable qualities of a small-molecular inhibitor, including high target potency and well-characterized selectivity, it is known for its inhibitory effect on lymphoid and hematopoietic tissues (35). JQ1 exhibits linear pharmacokinetics in mice with an oral bioavailability of approximately 50% and a half-life of 1 hour (*35, 36*). Due to the short half-life and potential off-target effects, this compound is only used in preclinical studies. Currently there are several BET inhibitors with better pharmacokinetic properties in clinical trials for various types of cancer (9). Recent trials have also focused on targeting specific BETs, including BRD4 inhibitors AZD5153 (NCT03205176), which was used in this study, and ABBC-744 (NCT03360006). Based on our results, BETs, in particular BRD4, regulate essential processes involved in SSc fibrosis. We not only revealed a novel anti-fibrotic mechanism involving calcium signaling by BET inhibition, but also highlighted the potential of epigenetic therapeutic strategies targeting BRD4 for SSc patients. These data, along with future studies focusing on additional molecular mechanisms underlying the role of BRD4 in SSc, should provide the framework that supports the use of more selective BRD4 inhibitors as a therapeutic option for SSc.

## Methods

### Bleomycin-induced skin fibrosis

The procedure to induce fibrosis by bleomycin in mice was published previously (2, 37). 100 μl of bleomycin (1 mg/ml) or PBS was injected subcutaneously into a single location on the shaved back of C57BL/6 mice once every day for 2 weeks. JQ1 (50 mg/kg, MedChemExpress) or vehicle control (20% DMSO/50% PBS/30% PEG) was given daily by oral gavage. In a separate experiment, BRD4 inhibitors AZD5153 (5mg/kg, Cayman), ARV825 (5mg/kg, MedChemExpress), or vehicle control (22% DMSO/48% PBS/30% PEG) were given daily intraperitonially. At the end of the experiment, fixed skin sections were stained with Masson’s trichrome. Dermal thickness was measured by analyzing the distance between the epidermal–dermal junction and the dermal–fat junction in three fields in two or more sections from each animal. Immunofluorescence was performed on sections using anti-αSMA (Abcam) or anti-F4/80 (Invitrogen) antibodies after antigen-retrieval. Collagen content in the skin was measured using the Hydroxyproline Kit (Abcam). All animal protocols were approved by the Institutional Animal Care & Use Committee at the University of Michigan.

### Patients and Controls

All patients recruited in this study met the ACR/EULAR criteria for the classification of SSc (38). Punch biopsies from the distal forearm of healthy volunteers and diffuse cutaneous (dc)SSc patients were obtained for fibroblast isolation. This study was approved by the University of Michigan Institutional Review Board. The demographics and clinical characteristics of the enrolled individuals are summarized in **Supplemental Table 2**.

### Cell culture

Punch biopsies obtained from healthy subjects and SSc patients were digested as previously described (2, 39). Briefly, biopsies were placed in 2.4 U/ml dispase overnight at 4°C. After removing the epidermal layers, the biopsies were transferred to 0.2% collagenase solution and incubated at 37°C for 45 min. The dissociated cells were then collected and cultured. After separating out the endothelial cells by magnetic bead selection, the resultant fibroblasts were maintained in RPMI supplemented with 10% fetal bovine serum (FBS) and antibiotics. Cells between passage 3 and 6 were used in all experiments.

### Cell treatment and transfection

Dermal fibroblasts from dcSSc patients were treated with 0.01-22 µM of JQ1 (Cayman), a pan-BET inhibitor, for 48 hours. Normal dermal fibroblasts were treated with TGFβ (10 ng/ml, Cell Signaling) and/or JQ1 (1 µM) for 72 hours to induce a myofibroblast phenotype. In a separate experiment, BRD2 inhibitor BIC1 (Cayman), BRD4 inhibitors AZD5153 or ARV825 (0.1-10 µM, Cayman), were used to treat dcSSc fibroblasts for 48 to 72 hours. BRD2, BRD3, or BRD4 knockdown in dcSSc dermal fibroblasts was performed using *BRD2* (200nM), *BRD3* (100nM), *or BRD4* (200nM) siRNA (ON-TARGETplus siRNA, Dharmacon). Scrambled siRNA (Dharmacon) was used as a control. The cells were transfected for 72 hours before downstream experiments were performed. *CAMK2A* in dcSSc fibroblasts was overexpressed using *CAMK2A* vectors from Origene. 0.1 μg/ml of *CAMK2A* vector was mixed with Lipofectamine 2000, then added to culture media. Inhibitors were added 24 hours after the transfection. PCMV6-XL6 vector (Origene) was used as a negativecontrol. To determine the effect of Ca^2+^ on myofibroblast transformation, dcSSc dermal fibroblasts were cultured in Ca^2+^-containing or Ca^2+^ -free DMEM media. In a separate experiment, dcSSc fibroblasts cultured in Ca^2+^-containing RPMI media were treated with BAPTA-AM (20 μM, Thermo Fisher) in the presence or absence of 1 μM JQ1 for 2 days.

### Western blots

Western blotting was performed following the protocol as described previously (39). Equal amount of protein was separate by SDS-PAGE, followed by transferring to a nitrocellulose membrane. After blocking, the blots were probed with antibodies against BRD2 (Abcam), BRD3 (Santa Cruz), BRD4 (Abcam), collagen I (Abcam), or αSMA (Abcam). For loading control, the blots were immunoblotted with antibodies against β-actin (Sigma) or GAPDH (Cell Signaling) as controls. Band quantification was performed using ImageJ (40).

### mRNA extraction and qRT-PCR

Total RNA was extracted using Direct-zol™ RNA MiniPrep Kit (Zymo Research) before converted to cDNA. Quantitative PCR was performed in a ViiA™ 7 Real-Time PCR System.

### Cell proliferation assays

Proliferation of dermal fibroblasts was measured using the BrdU Cell Proliferation Assay Kit (BioVision). After 4 hours of BrdU incubation, cells were fixed before adding antibodies and substrate. The absorbance at 450 nm was measured. In a separate experiment, the IncuCyte® Live-Cell Imaging System was used to monitor cell proliferation. Cells were seeded and allowed to grow overnight.After adding different treatments cells were monitored by IncuCyte® up to 5 days. Cell counts were analyzed by the IncuCyte® S3 Analysis software.

### Gel contraction and cell migration assays

The Cell Contraction Assay kit (Cell Biolabs) was used for gel contraction (39). After treatment, cells were suspended and mixed with collagen solution. Solidified gels were lifted after 24 hours and the areas of the gels were quantified using ImageJ (40). To evaluate the effect of JQ1 on cell migration, we performed a scratch wound assay using the IncuCyte® Live-Cell Imaging System. Cell migration was monitored by IncuCyte® up to 7 days.

### Detection of apoptosis

To assess the effect of JQ1 on cell apoptosis, cells were plated in 96 well plates and treated with JQ1 in the presence of IncuCyte® Caspase-3/7 Apoptosis Reagent. The apoptotic cells were quantified by measuring cells with green fluorescent staining. The data were presented as green fluorescent cell number normalized to total cell count.

### Cell cycle analysis

Fibroblasts were treated with 1 µM JQ1 for 48 hours, while the non-treated dcSSc fibroblasts were used as control. Both the JQ1- and non-treated cells were first synchronized using the double thymidine block method before treatment (41). Cells were then harvested and stained with 7-AAD viability dye and Vybrant DyeCycle Violet (Invitrogen) according to the manufacture’s protocol. The stained cells were analyzed using a BD FACSCanto II flow cytometer. Cell cycle analysis was performed on gated 7-AAD-live cells using Flowjo 7.0.

### RNA sequencing and analysis

RNA-seq was performed on mRNA extracted from control- or JQ1 (1 μM)-treated dcSSc fibroblasts. Samples with an RNA-integrity number of more than 7 were subjected to library preparation and sequencing to 151 paired-end cycles on the NovaSeq-6000 platform (Illumina), resulting in approximately 35 million reads/sample. Differential gene expression analysis was performed using DESeq2 (42), using a negative binomial generalized linear model (thresholds: linear fold change > 1.5 or < -1.5, Benjamini-Hochberg FDR (Padj) <0.05). Plots were generated using variations of DESeq2 plotting functions and other packages with R version 3.3.3. Functional analysis, including candidate pathways activated or inhibited in comparison(s) and GO-term enrichments, was performed using iPathway Guide (Advaita)(43). Additional functional enrichment analyses to generate networks for visualization were performed using ClueGO (v2.5.7)/CluePedia (v1.5.7)(44, 45) and Cytoscape (v3.8.0)(46).

### Intracellular Ca^2+^ measurement

Dermal fibroblasts from dcSSc patients were treated with or without 1 µM JQ1, AZD5153, ARV825, or BIC1 for 48 hours before being collected for Ca^2+^ measurement by flow cytometry. Detached cells (1×10^7^/mL) were labeled in loading buffer (HBSS with 1% FBS, 1 mM CaCl_2_ and 1 mM MgCl_2_) containing 1 μM Fluo-4 AM (Thermo Fisher) with 0.02% Pluronic F-127 at 37 °C for 30 min with gentle agitation every 10 min. The labeled cells were washed twice with loading buffer containing 0.1 mM sulfinpyrazone, and resuspended in the same loading buffer with sulfinpyrazone at 1×10^6^/mL, incubated at 37°C for 30 min before basal fluorescence of Fluo-4 were measured on a BD FACSCanto II flowcytometer. Mean fluorescent intensity of unlabeled cells was subtracted from the mean fluorescent intensity of the labeled cells.

### Statistics

Normality test was conducted to determine whether the data is normally distributed or skewed. To determine the differences between groups, unpaired t test, Mann–Whitney U test, paired t-test, Wilcoxon test, one-way ANOVA with Sidak test, Kruskal–Wallis test with Dunn’s test, or two-way ANOVA with Dunnett’s test were performed using GraphPad Prism version 8 (GraphPad Software, Inc). P values of less than 0.05 were considered statistically significant. Results were expressed as mean ± SD.

### Study Approval

This study was approved by the University of Michigan Institutional Review Board. Written informed consent was received from participants prior to inclusion in the study. All animal protocols were approved for use from the University of Michigan the Institutional Animal Care & Use Committee.

## Supporting information

All supplemental files

## Author contributions

All authors participated in the interpretation of study results, and in the drafting, critical revision and approval of the final version of the manuscript. PT, AHS, DF, YMD, and DK contributed to study conception and/or design. SV, MG, MNM, WDB, SM, PJP, MA, PLC, MAA, JHR, QW, EM, DMR, JLH, and PT contributed to the acquisition of study results. SV, PJP, MA, QW, and PT contributed to the analysis of study results.

## Acknowledgements

This work was supported by the funds from the American Autoimmune Related Disease Foundation and the Edward T. and Ellen K. Dryer Early Career Professorship (Tsou), Chugai, PCORI, Novartis, Sanofi-Genzyme, and Genentech (Mao-Draayer), National Institute of Arthritis and Musculoskeletal and Skin Diseases grant number K24AR063120 (Khanna) and R01AR070148 (Sawalha), and National Institute of Allergy and Infectious Diseases grant number UM1-AI110557-05 and UM1-AI144298-01 (Mao-Draayer), and R01AI097134 (Sawalha).

## Data availability

All data relevant to the study are included in the article. The RNA-sequencing data is deposited to GEO (GSE186961).

