## Supplementary material for "Inhibition of histone readers bromodomain extra-terminal proteins alleviates skin fibrosis in experimental models of scleroderma": All supplemental files

**Supplemental Figure 1. Inhibition of BETs by JQ1 affects genes involved in fibrosis in dcSSc fibroblasts.** At doses between 0.01-22 μM, JQ1 significantly downregulated *ACTA2*, *COL1A1*, *CTGF* and *BRD4* in dcSSc fibroblasts, and upregulated *MMP1*, *TGFB1*, and *BRD2*. JQ1 did not affect *TIMP1* and *BRD3* expression. n=number of patients. Results are expressed as mean +/- SD and p<0.05 was considered significant.

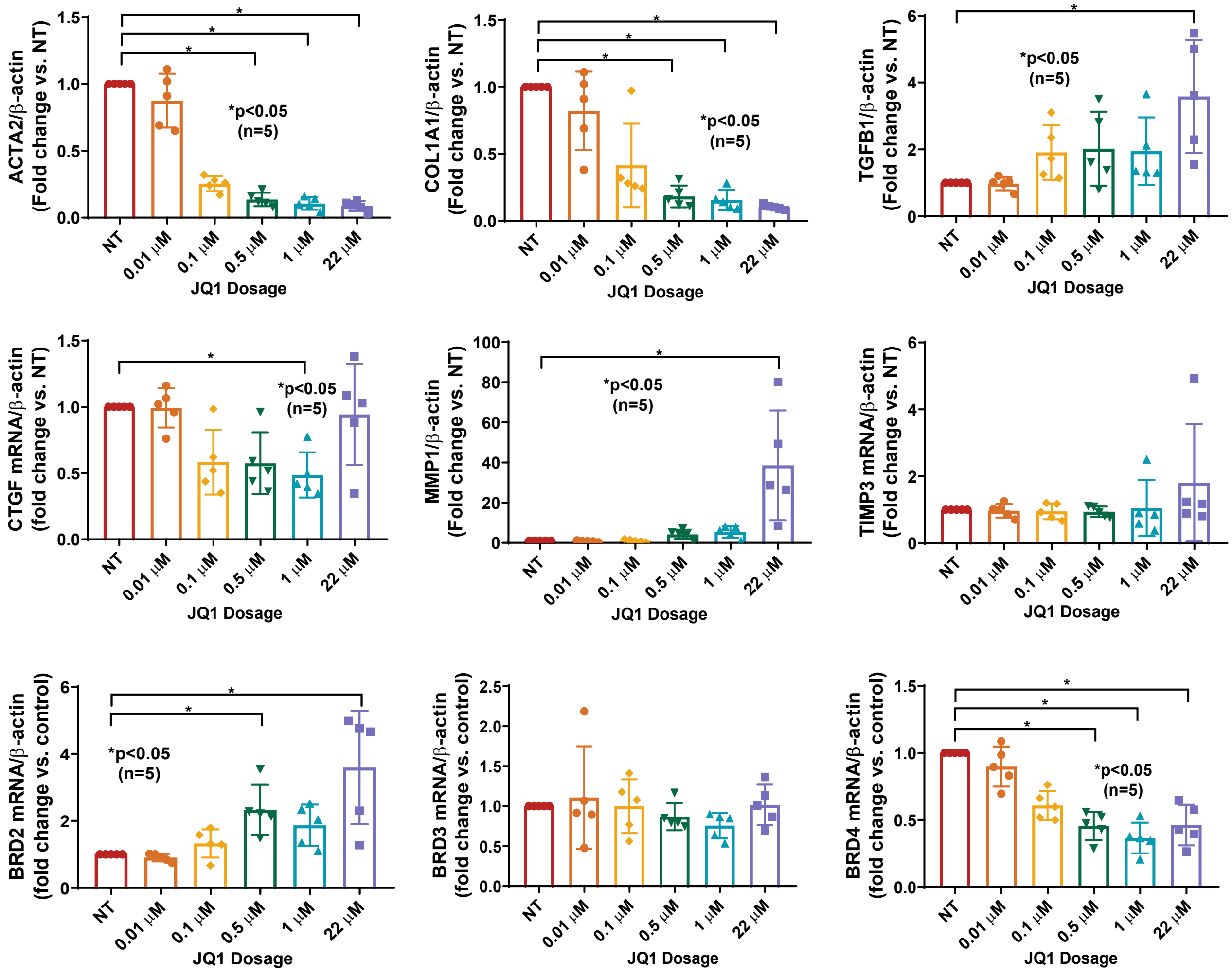

**Supplemental Figure 2. Inhibition of proliferation by JQ1 is mediated by apoptosis and cell cycle arrest in dcSSc fibroblast. (A)** JQ1 induced significant cell apoptosis in dcSSc fibroblasts. Apoptosis was measured by a green fluorescence dye that couples with activated caspase 3/7 in cells. Apoptotic cell count and total cell numbers were monitored by IncuCyte® Live-cell imaging. Data was presented as number of apoptotic cells normalized by total cell count. Representative result from 3 patient lines. **(B)** JQ1 (1μM) induced cell cycle arrest in dcSSc fibroblasts; it induced cell accumulation in G0/G1 phase and a decrease in S phase. n=number of patients. Results are expressed as mean +/- SD and p<0.05 was considered significant.

A

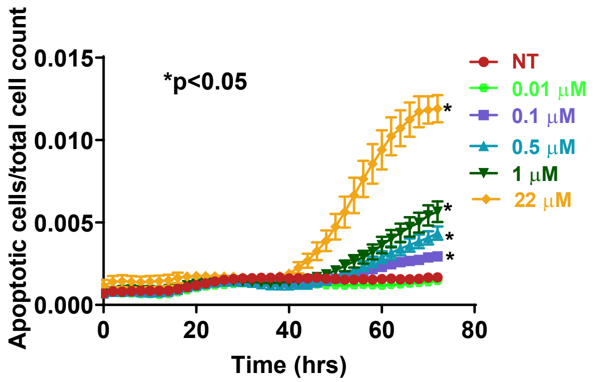


B

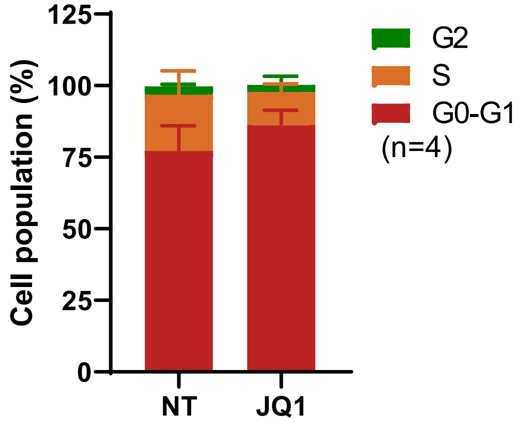

**Supplemental Figure 3. Heat map of top 500 genes significantly altered by JQ1 (pink) vs. control (blue).** Scleroderma (SSc) dermal fibroblasts were treated with 1μM JQ1 for 48 hours and RNA was extracted for RNA-seq analysis. Each column represents data from a SSc patient (n=6 pairs).

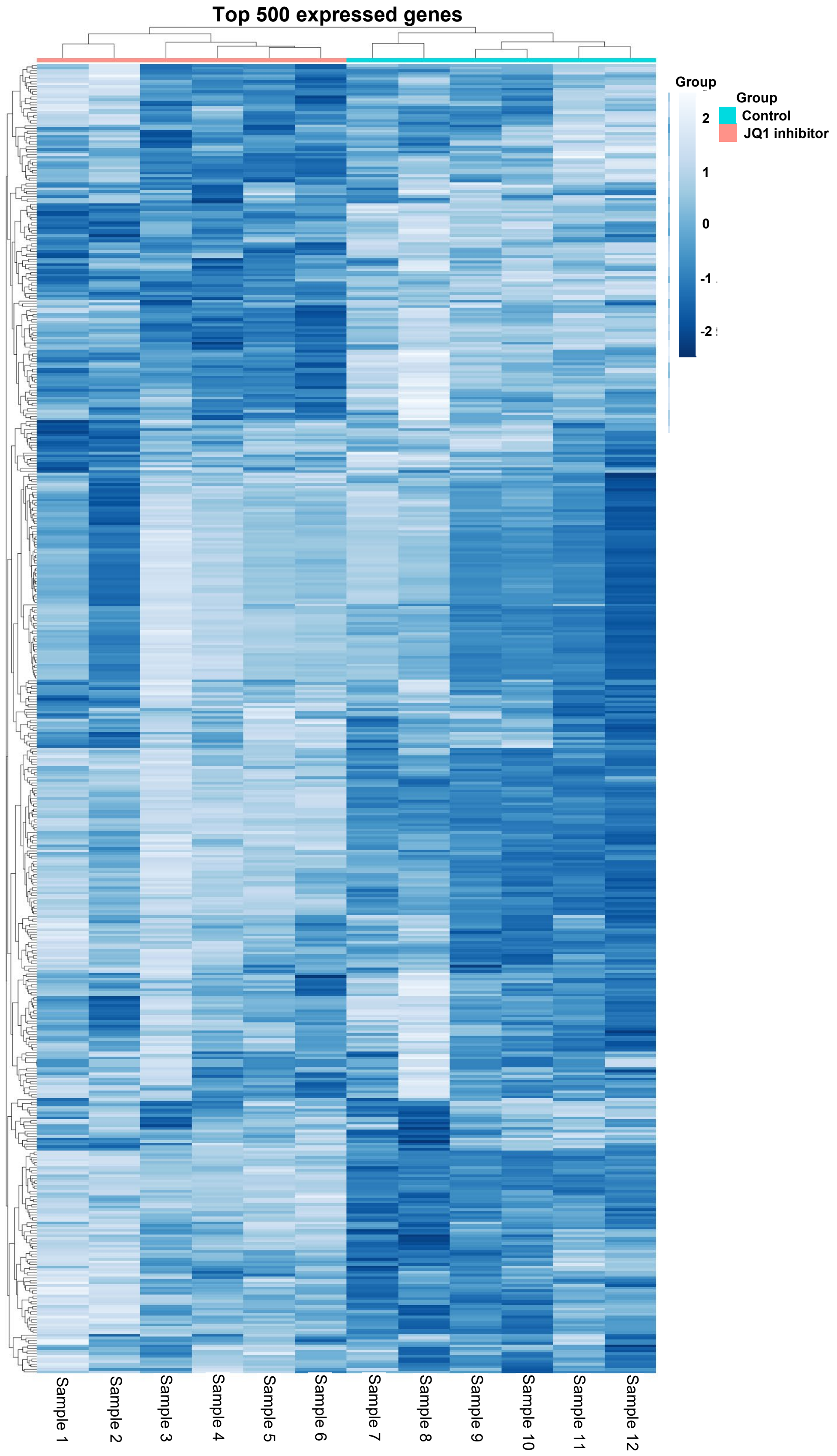

**Supplemental Figure 4. Differentially expressed genes after JQ1 treatment in dcSSc fibroblasts from RNA-seq analysis showed significant downregulation of *ACTA2*, *COL1A1* (blue bars), and upregulation of *TGFB1* and *BRD2* (red bars).** Genes that are not significant after correction for multiple testing are represented by unfilled bars.

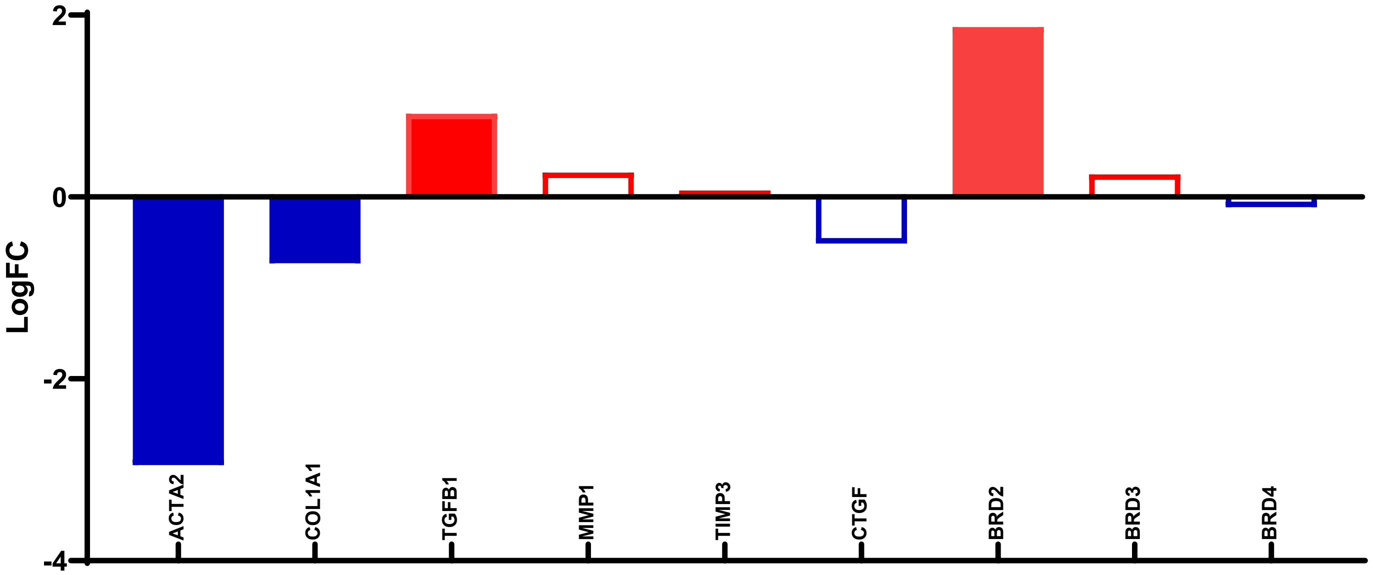

**Supplemental Figure 5. Functional grouping and subcellular localization of the differentially expressed genes after JQ1 treatment mapped to subcellular regions (labeled on the right).** This analysis was done using the Cerebral layout function in ClueGO. The node size shows the term significance; Node size is proportional to significance.

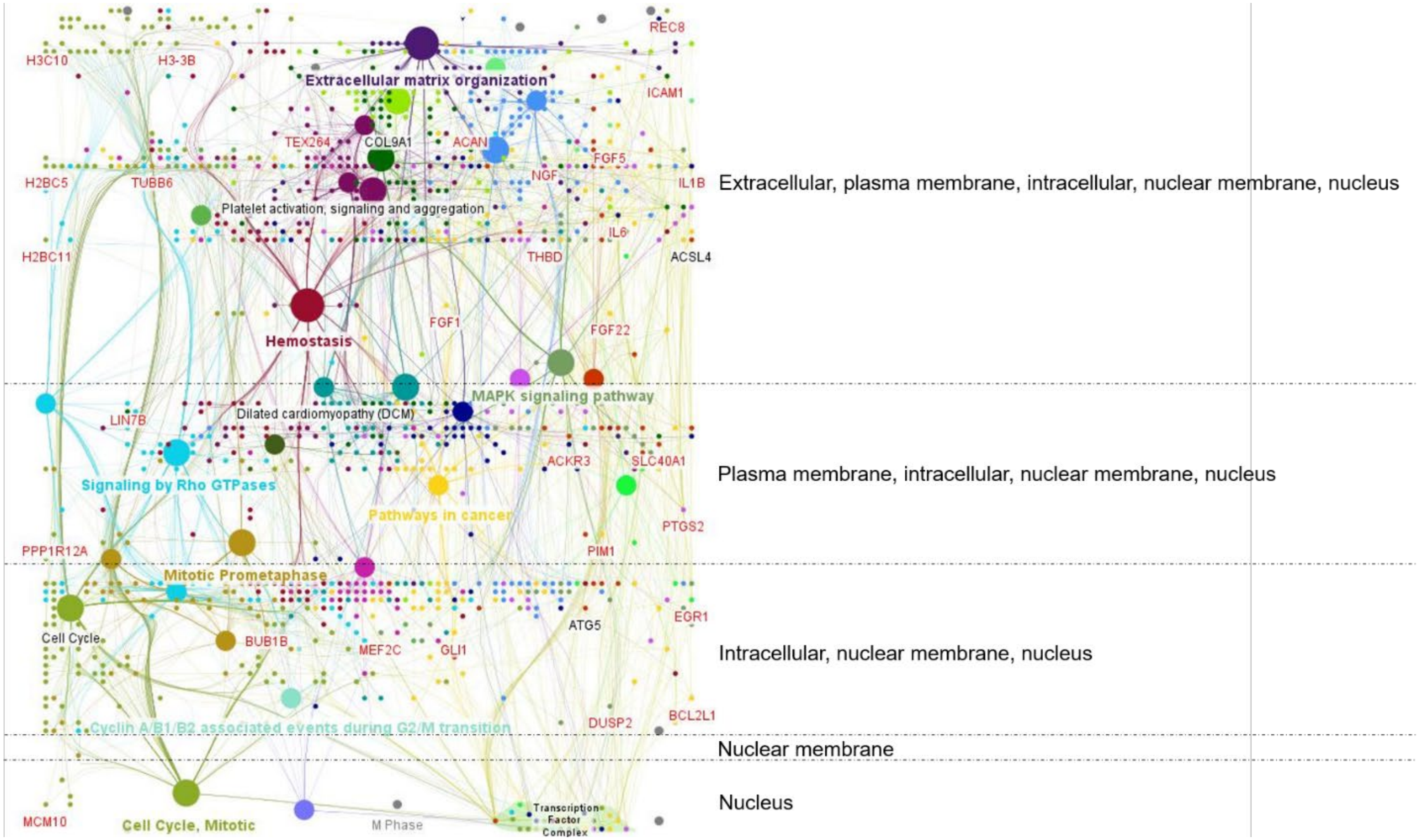

**Supplemental Figure 6. Differential expressed genes after JQ1 treatment in SSc fibroblasts enriched in ECM-Receptor Interaction (KEGG: 04512). (A)** The pathway diagram is overlaid with the expression changes of each gene. The most downregulated expression is shown in dark blue, while the most upregulated expression is in dark red. The legend describes the values on the gradient. The genes enriched in each pathway are also listed. **(B)** All the differentially expressed genes in the KEGG: 04512 ECM-Receptor Interaction are ranked based on their absolute value of log fold change. Upregulated genes are shown in red, downregulated genes are shown in blue.

**A**

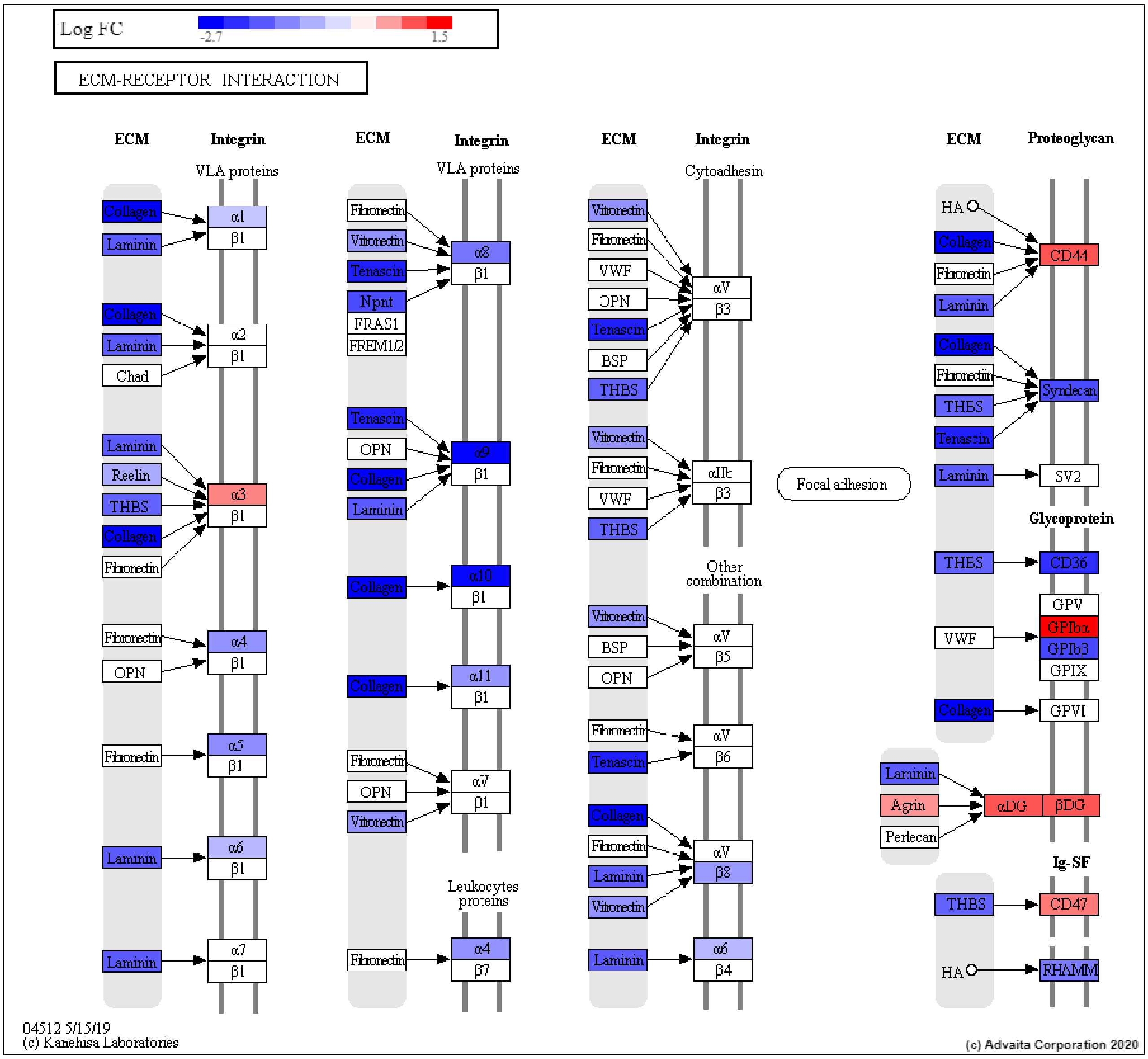

B

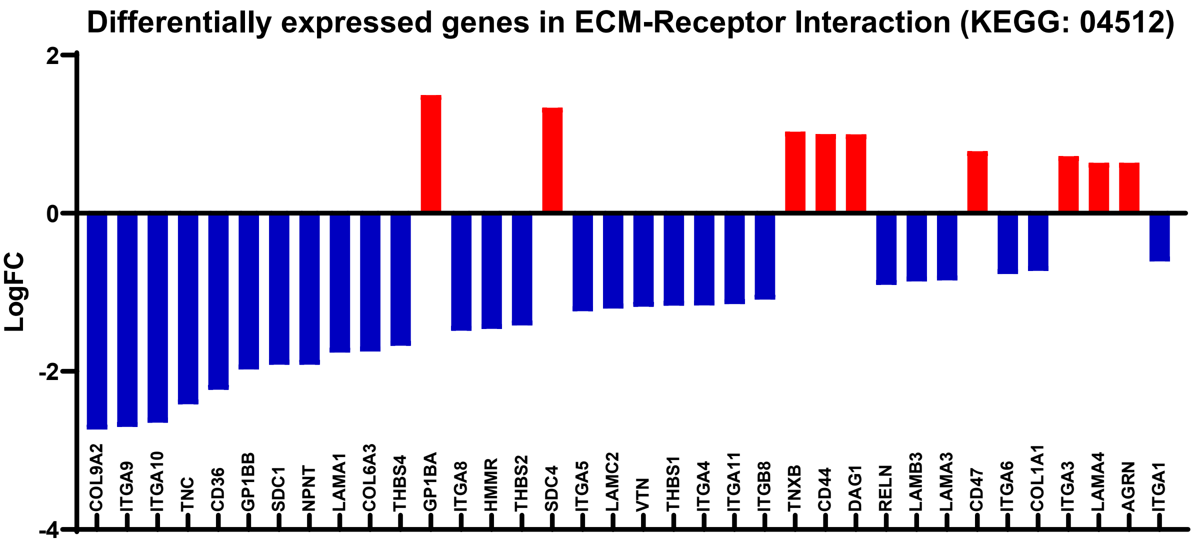

Differentially expressed genes in ECM-Receptor Interaction (KEGG: 04512)
LogFC
2
0
-2
-4
COL9A2
ITGA9
ITGA10
TNC
CD36
GP1BB
SDC1
NPNT
LAMA1
COL6A3
THBS4
GP1BA
ITGA8
HMMR
THBS2
SDC4
ITGA5
LAMC2
VTN
THBS1
ITGA4
ITGA11
ITGB8
TNXB
CD44
DAG1
RELN
LAMB3
LAMA3
CD47
ITGA6
COL1A1
ITGA3
LAMA4
AGRN
ITGA1

**Supplemental Figure 7. Predicted upstream regulator analysis using the differentially expressed genes from RNA-seq after JQ1 treatment predicted JQ1 as the most overly abundant, as 1521 genes from our transcriptomic analysis overlapped with reported JQ1-target genes. (A)** Present (overly abundant) p-value vs zscore p-value plot: the significance of JQ1 (yellow circle) is plotted on two axes, with negative log of $P_z$ on x-axis and negative log of $P_{pres}$ on y-axis. The size of the dot represents the relative number of consistent differentially expressed genes comparing genes from our RNA-seq result and various upstream regulators. The yellow circle shows that 1521 genes from our RNA-seq data overlapped with reported JQ1-target genes. **(B)** Volcano plot: The 1521 differentially expressed genes from our RNA-seq data overlapped with reported JQ1-genes are shown here. The target genes are represented in terms of their measured expression change (x-axis) and the significance of the change (y-axis). The dotted lines represent the thresholds used to select the differentially expressed genes: 0.585 for expression change and 0.05 for significance.

**A**

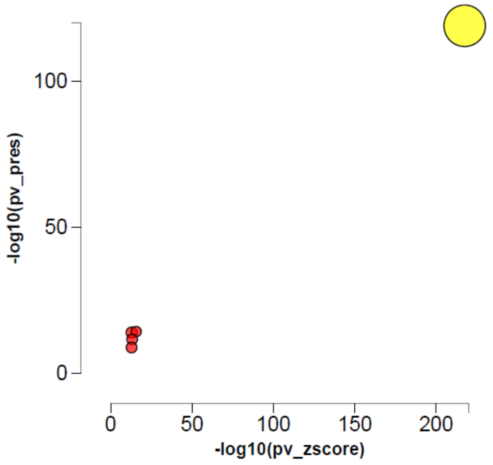


**B**

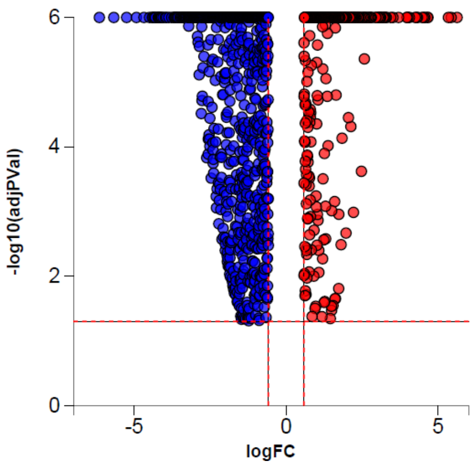

**Supplemental Figure 8. The expression of BET proteins is not altered in dcSSc compared to normal fibroblasts. (A)** At the mRNA levels, *BRD2* and *BRD3* were similar in normal and dcSSc fibroblasts, while *BRD4* was significantly upregulated in dcSSc fibroblasts. **(B)** At the protein level, BRD2, BRD3, and BRD4 showed variable levels in dcSSc, however there was no difference between normal and dcSSc fibroblasts when the bands were quantified. n=number of patients. Results are expressed as mean +/- SD and p<0.05 was considered significant.

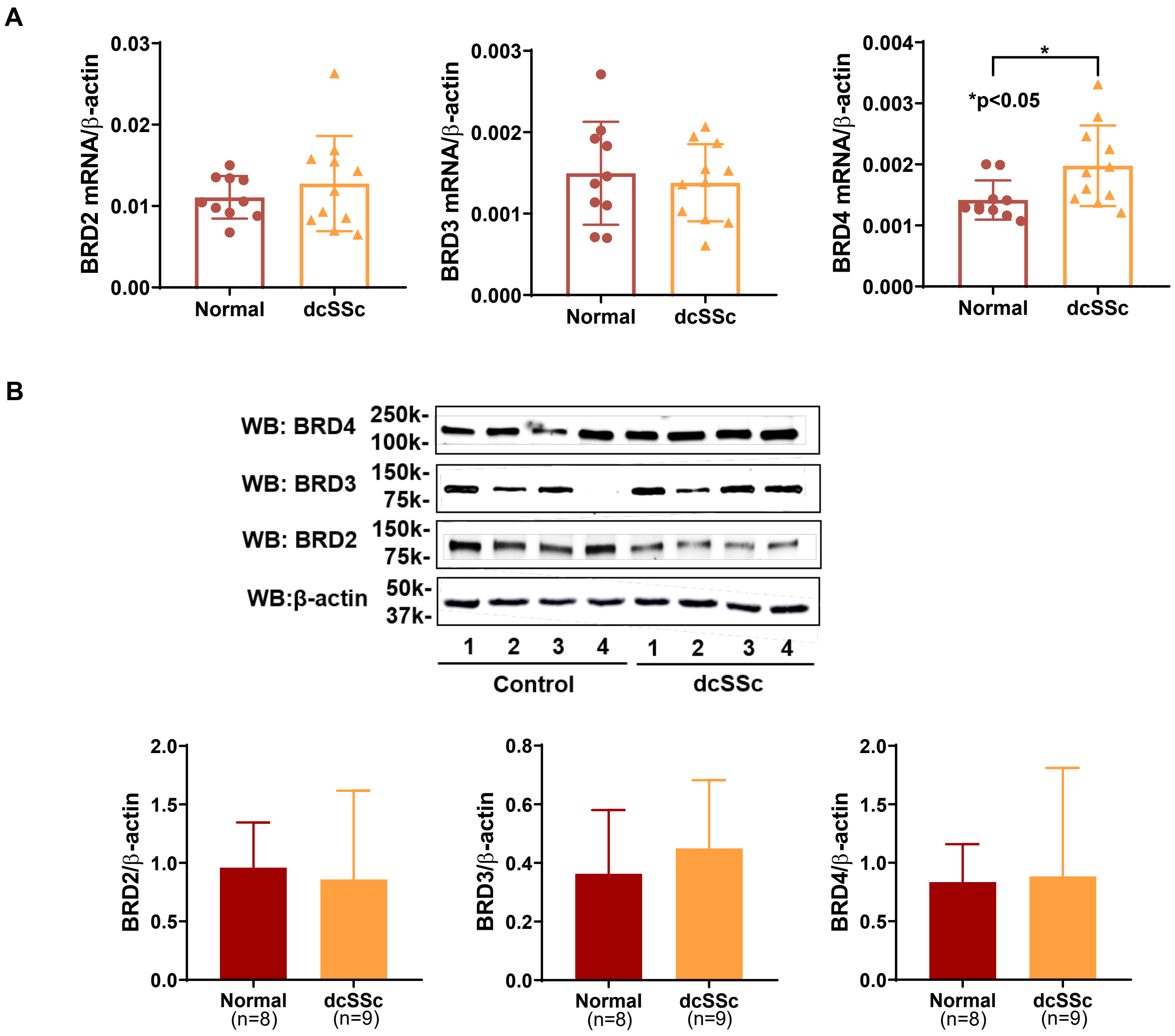

**Supplemental Figure 9. Genome browser tracks of the *ACTA2* and *COL1A1* loci generated in primary human dermal fibroblasts.** Tracks for RNA-seq, DNA methylation, DNase I hypersensitivity, and ChIP-seq data for acetylated histone marks were extracted from the ENCODE database.

***ACTA2***

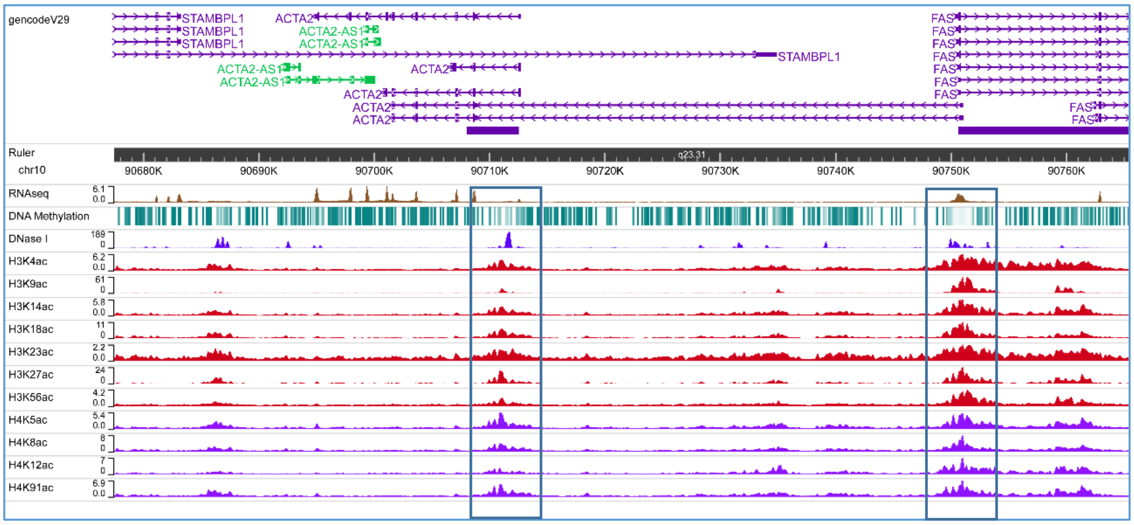


***COL1A1***

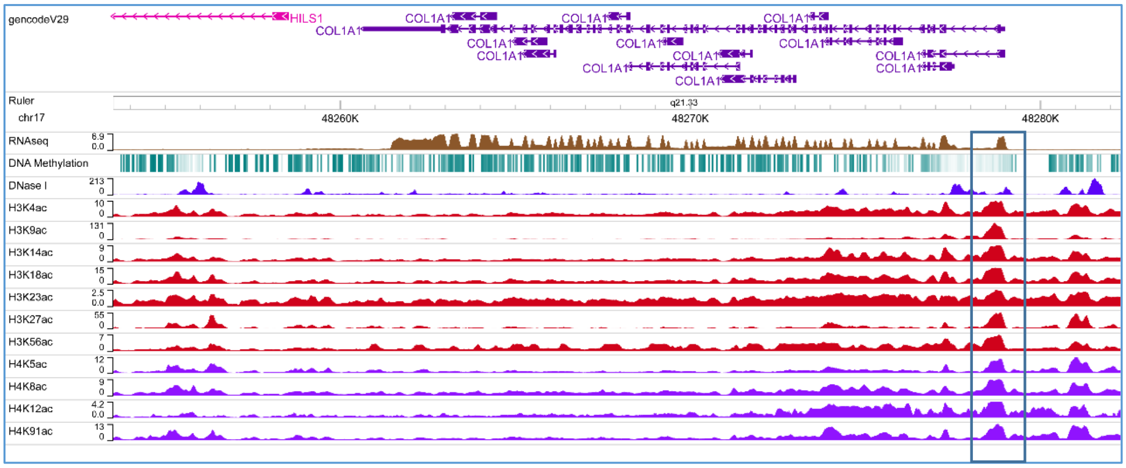

**Supplemental Table 1:** Pathway enrichment analysis of differentially expressed genes in dcSSc fibroblasts after JQ1 treatment. DEG: differentially expressed genes

| KEGG Pathway | No. DEG | Total No. of Genes | Adjusted P Value |
|---|---|---|---|
| Calcium signaling pathway | 59 | 140 | 0.0014 |
| Cytokine-cytokine receptor interaction | 71 | 160 | 0.0014 |
| MAPK signaling pathway | 103 | 259 | 0.0014 |
| Rap1 signaling pathway | 72 | 175 | 0.0014 |
| Hypertrophic cardiomyopathy (HCM) | 38 | 70 | 0.0014 |
| Malaria | 21 | 29 | 0.0015 |
| Systemic lupus erythematosus | 42 | 78 | 0.0029 |
| Neuroactive ligand-receptor interaction | 65 | 137 | 0.0054 |
| Pathways in cancer | 161 | 441 | 0.0099 |
| Biosynthesis of amino acids | 32 | 62 | 0.0099 |
| Vascular smooth muscle contraction | 43 | 97 | 0.0099 |
| Inflammatory bowel disease (IBD) | 24 | 39 | 0.013 |
| Metabolic pathways | 401 | 1162 | 0.013 |
| Dilated cardiomyopathy (DCM) | 38 | 73 | 0.014 |
| Arginine and proline metabolism | 21 | 38 | 0.024 |
| Ferroptosis | 21 | 35 | 0.025 |
| MicroRNAs in cancer | 59 | 139 | 0.026 |
| Glutathione metabolism | 24 | 46 | 0.027 |
| Inflammatory mediator regulation of TRP channels | 37 | 80 | 0.028 |
| Salivary secretion | 23 | 53 | 0.028 |
| Cell adhesion molecules (CAMs) | 44 | 99 | 0.028 |
| PI3K-Akt signaling pathway | 98 | 282 | 0.030 |
| ECM-receptor interaction | 36 | 74 | 0.040 |
| Hematopoietic cell lineage | 26 | 53 | 0.040 |

**Supplemental Table 2:** SSc patients and healthy controls characteristics.

| | dcSSc (n=44) | Healthy volunteers (n=21) |
|---|---|---|
| Age (years) | 56.5 ± 2.1[a] | 56.4 ± 3.3 |
| Sex | F31/M13 | F11/M10 |
| Disease duration (years) | 3.1 ± 0.5 | N.A. |
| Modified Rodnan Skin Score | 18.2 ± 1.7 | N.A. |
| Raynaud's phenomenon | 41 | N.A. |
| Early disease (< 5yrs) | 40 | N.A. |
| Interstitial lung disease | 28 | N.A. |
| Pulmonary arterial hypertension | 9 | N.A. |
| Immunosuppressive | 36 | N.A. |

[a]Mean ± SEM

[b]N.A. = Not applicable
